# Retinoic Acid Boosts HIV-1 Replication in Macrophages *via* CCR5/SAMHD1-Dependent and mTOR-Modulated Mechanisms

**DOI:** 10.1101/2023.10.05.561142

**Authors:** Jonathan Dias, Amélie Cattin, Jean-Philippe Goulet, Augustine Fert, Laurence Raymond Marchand, Tomas Raul Wiche Salinas, Christ-Dominique Ngassaki Yoka, Etiene Moreira Gabriel, Edwin Ramon Caballero, Jean-Pierre Routy, Petronela Ancuta

## Abstract

The intestinal environment facilitates HIV-1 infection *via* mechanisms involving the gut-homing elixir retinoic acid (RA), which transcriptionally reprograms CD4^+^ T-cells for increased HIV-1 permissiveness. Consistently, colon-infiltrating CD4^+^ T-cells carry replication-competent viral reservoirs in people living with HIV-1 (PLWH) receiving antiretroviral therapy (ART). Intriguingly, integrative infection in colon macrophages, a pool replenished by circulating monocytes, represents a rare event in ART-treated PLWH, thus questioning on HIV-1 permissiveness in gut-resident macrophages. Here, we demonstrate that RA significantly boosts R5 but not X4 HIV-1 replication in monocyte-derived macrophages (MDMs). RNA-Sequencing, Gene Set Variation Analysis, and HIV interactor NCBI database interrogation, revealed RA-mediated transcriptional reprogramming associated with metabolic/inflammatory processes and HIV-1 resistance/dependency factors. Functional validations pointed to mechanisms of RA action, including CCR5 upregulation and SAMHD1 phosphorylation under the control of mTOR. These results support a model in which intestinal MDM contribute to viral replication/dissemination before ART and upon treatment interruption in mTOR-sensitive manner.

**Figure 1:**
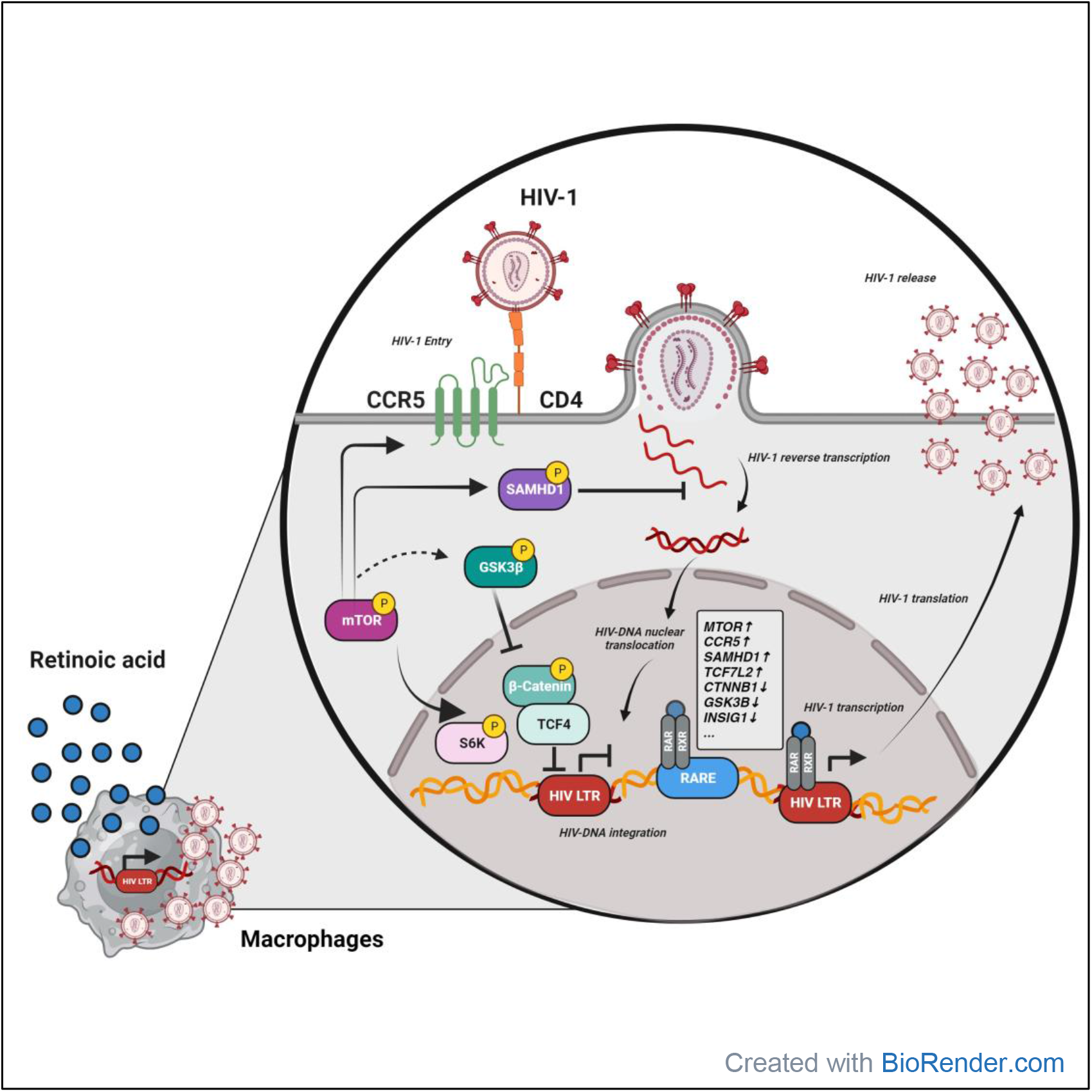

## HIGHLIGHTS AND ETOC BLURB

### Highlights

- Retinoic acid (RA) boosts R5 but not X4 HIV-1 replication in macrophages
- RA facilitates HIV-1 replication *via* entry and post-entry mechanisms
- RA activates mTOR, which blunts SAMHD1-mediated HIV-1 restriction
- mTOR inhibitors decrease CCR5 expression and increases SAMHD1 activity

### eTOC Blurb

Dias *et al.* investigated that effect of RA, the gut-homing elixir, on macrophages. RA boosted permissiveness to R5 HIV-1 strains by upregulating CCR5 expression and the efficacy of post-entry steps, before/after HIV-1 integration. RNA-Sequencing and functional validations identified HIV interactors as direct RA targets and revealed SAMHD1-dependent and mTOR-modulated mechanisms.

## INTRODUCTION

Despite the efficacy of antiretroviral therapy (ART) in controlling HIV-1 replication to undetectable plasma levels, ART does not eradicate HIV-1^1-3^. The major barrier to HIV-1 cure is the persistence of viral reservoirs (VRs) carrying integrated HIV-DNA during ART; this explains the rapid viral rebound upon treatment interruption and imposes a life-long treatment in people living with HIV (PLWH)^2-5^. Recent studies on autopsy tissues from ART-treated PLWH documented the presence of intact VRs in the majority of tissues investigated^6-8^. The persistence of VRs is well-documented in long-lived memory CD4^+^ T cells and multiple investigation efforts were made to purge VRs in this cellular compartment^3,9^. In addition, other immune cells such as macrophages residing in deep tissues may represent sanctuaries for VRs during ART, but remain poorly investigated due to restricted biological sample accessibility^10-17^. In addition to their role as HIV-1 integrative infection sites, macrophages also exhibit the ability to disseminate HIV-1 to CD4^+^ T-cells^12,18,19^.

Pioneering studies in the pre-ART era documented HIV-1 infection of macrophages in tissues of PLWH^20,21^. In line with this observation, macrophages express the HIV-1 receptor CD4 and co-receptors CCR5 and CXCR4 and support productive HIV-1 infection *in vitro*^22-27^. Other mechanisms of infection in macrophages include cell-to-cell transmission, fusion with, or phagocytosis of infected CD4^+^ T-cells^28-31^. Of note, the expression of *do-not-eat-me* signals is modulated by HIV-1 in CD4^+^ T-cells, thus facilitating phagocytosis by macrophages for subsequent productive infection^32^. In addition to macrophage permissiveness to HIV-1 infection *in vivo* and *in vitro*, studies in ART-treated PLWH provided evidence that macrophages isolated from liver^33^ brain^34,35^, broncho alveolar lavage^36-38^, duodenum^39^, testis^40^, and most recently urethra^41^ carry HIV-1. Similarly, studies performed in humanized mice models, especially in myeloid only models^42,43^, as well as SIV infection model^44-46^ pointed to macrophages as important sites of viral infection/persistence. Finally, by their intrinsic resistance to apoptosis and CD8^+^ T-cell- and NK cell-mediated killing^47-49^, HIV-infected macrophages escape from immunological pressure.

The persistence of VRs in macrophages depends on the long-term survival potential and self-renewal capacity of such cells in ART-treated PLWH, features that vary among macrophage subsets relative to their ontogeny. Tissue-resident macrophages (TRM) from the brain (microglia), liver (Kupffer cells), lungs (alveolar macrophages) and epidermis (Langerhans cells) derive from embryonic/fetal precursors during intrauterine development and represent long-lived self-renewing TRM^50,51^. In contrast, fractions of macrophages from the heart, pancreas, intestine, and dermis derive from bone marrow-derived monocytes and represent short-lived macrophages^50,51^. Advances in the past decade demonstrated that long-lived TRM exist in all tissues and their role is mainly in tissue remodeling/homeostasis^50,51^. This pool of self-renewing TRM progenitors suffer attrition with ageing, thus allowing a niche for pro-inflammatory short-lived macrophages derived from circulating monocytes to infiltrate^52^. This is the case for cardiac macrophages, as demonstrated by mouse^53^ and human studies^54^. Whether macrophages with distinct ontogeny residing in specific tissues differentially support HIV-1 replication and/or VR persistence remains a field of open investigations.

The intestine is an anatomic site containing the vast majority of immune cells, which are preferentially targeted by HIV-1 for infection^55-58^ and VR persistence during ART^59-62^. The intestinal environment is rich in the gut-homing “*elixir*” retinoic acid (RA)^63-65^, a metabolic derivative of vitamin A, produced by mucosal dendritic cells (DCs) expressing retinaldehyde dehydrogenase (RALDH) activity^66,67^. RA enter the cells via the STRA6 receptor and is transported by CRABP2 to the RA receptor alpha (RARA), which forms a heterodimer with RXR^68^. The RARA/RXR heterodimer undergoes nuclear translocation and binds on RA responsive elements (RARE) on the promoter of specific genes^68^, including the gut-homing integrin beta 7 (ITGB7) and the chemokine receptor CCR9^63^. Importantly, the HIV-1 LTR also contains RARE^69^, indicative of a direct role of RA on HIV-1 transcription. Our group demonstrated that RA was preferentially produced by CD16^+^ monocyte-derived DC upon interaction with specific components of the microbiota^70,71^ and that RA transcriptionally reprogramed CD4^+^ T-cells for increased HIV-1 replication/outgrowth^60,72-74^. Consistently, our group and others demonstrated that colon-infiltrating CD4^+^ T-cells carry VRs in PLWH receiving ART^60-62,75,76^. Intriguingly, our studies revealed that integrative infection in colon macrophages, a pool constantly replenished by circulating monocytes^50,51^, represents a rare event in ART-treated PLWH^76^. These findings raised questions on the effect of RA on HIV-1 permissiveness in macrophages from this anatomic site.

In this manuscript, we explored the effects of RA on HIV-1 replication in monocyte-derived macrophages (MDMs) and sought to identify the underlying molecular mechanisms of action. Our results demonstrate that RA increased CCR5 but decreased CXCR4 expression and rendered MDMs highly permissive to R5 HIV-1 replication, with no influence on X4 HIV-1 strains. These findings point to a putative role for RA in the selection of R5 HIV-1 strains. Single-round infection with VSV-G pseudotyped HIV-1 indicated that RA also acted at post-entry levels by mechanisms involving efficient HIV-1 reverse transcription and transcription/translation. RNA- Sequencing using the Illumina technology revealed a profound RA-mediated transcriptional reprogramming coinciding with the modulation of mTOR/S6K and Wnt/β-catenin/TCF4 signaling pathways. Functional validations revealed that RA induced CCR5 expression and reduced the HIV-1 restriction activity of SAMHD1 in mTOR-dependent manner. These results support a model in which macrophages in a RA-rich environment, such as the intestine, contribute to primary R5 HIV-1 infection before ART initiation, as well as viral rebound upon ART interruption. These results also point to mTOR as a therapeutic target to counteract the effects of RA on MDMs. Finally, the rarity of VR detection in colon-infiltrating macrophages of ART-treated PLWH ^76^ is likely not due to their resistance to HIV-1 infection, but rather explained by the rapid turnover of the intestinal macrophage pool that is constantly replenished by monocytes *in vivo*.

## RESULTS

### ATRA increases CCR5-tropic HIV-1 replication in macrophages

To investigate the effects of RA on HIV-1 replication in macrophages, MDMs were obtained by culturing highly pure monocytes, isolated by negative selection from PBMC of HIV-uninfected participants, in the presence of M-CSF (Figure 1A; Supplemental Figure 1A-C). MDMs were generated in the presence (ATRA-MDMs) or the absence (DMSO-MDMs) to *all-trans* RA (ATRA). Prior HIV-1 exposure, MDMs were analyzed by flow cytometry for the expression of the HIV-1 receptor CD4 and co-receptors CCR5 and CXCR4 ^77,78^ (Figure 1B). A significant increase in CCR5 and a decrease in CD4 and CXCR4 mean fluorescence intensity (MFI) expression were observed in ATRA-MDMs *versus* DMSO-MDMs, indicative that ATRA may facilitate CCR5-mediated HIV entry in MDMs (Figure 1C). To test this possibility, MDMs were exposed to the replication-competent CCR5-tropic (R5) HIV-1 strains, HIV_NL4.3BaL_ or transmitted founder (T/F) HIV_THRO_. In the case of HIV_NL4.3BaL_, ATRA significantly increased HIV-DNA integration at day 3 post-infection (n=8; p=0.0078) (Figure 1D). Consistently, HIV- p24 levels in cell-culture supernatants, indicative of productive infection, were increased by ATRA, with statistically significant differences observed at day 9 post-infection (n=8; p=0.0078) (Figure 1E). Similar results were observed with HIV_THRO_, where integrated HIV-DNA levels at day 3 post-infection (n=13; p=0.0002) (Figure 1F), as well as HIV-p24 levels in cell-culture supernatants collected at day 9 post-infection, were significantly increased by ATRA (n=8; p=0.0078) (Figure 1G).

**Figure 1:**
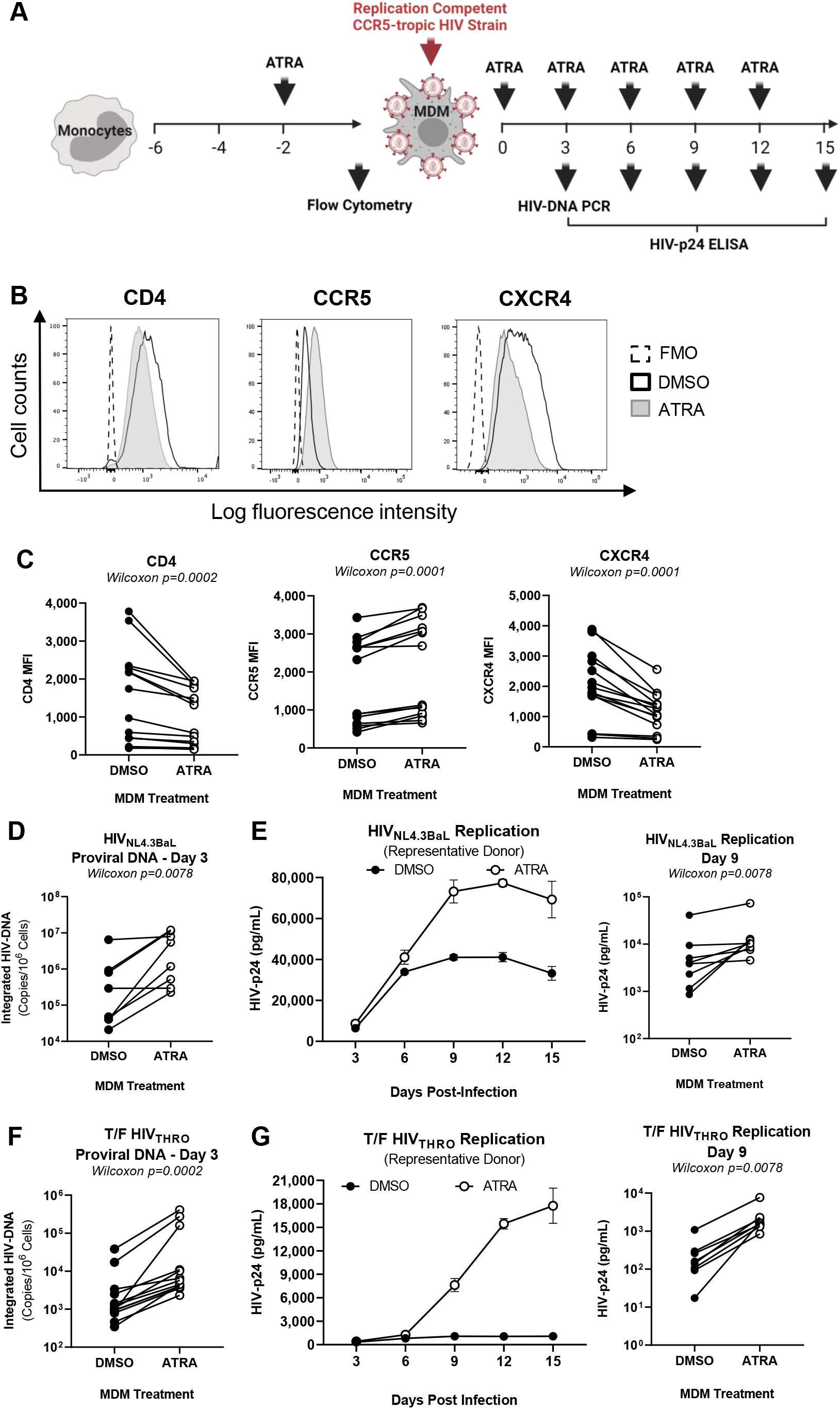
ATRA increases CCR5 expression and R5 HIV-1 replication in macrophages. Shown is the experimental flowchart **(A)**. Briefly, monocyte-derived macrophages (MDMs) were obtained by culturing monocytes in media containing M-CSF (20 ng/mL) for six days. MDMs were exposed (ATRA-MDMs) or not to ATRA (DMSO-MDMs) (10 nM) before and after HIV-1 exposure. **(B-C)** Prior to HIV-1 exposure, MDMs were analyzed by flow cytometry upon staining with CD4, CCR5 and CXCR4 antibodies. Shown are histograms for CD4, CCR5 and CXCR4 expression on MDMs from one representative donor **(B)**, as well as statistical analysis of CD4 **(left panel)**, CCR5 **(middle panel)** and CXCR4 **(right panel)** MFI expression on MDMs from n=14 participants. In parallel, MDMs were exposed to replication-competent CCR5-tropic HIV-1 strain (R5, HIV_NL4.3BaL_ or transmitted founder (T/F) HIV_THRO_) and cultured in media containing M-CSF in the presence/absence of ATRA for 15 additional days. Cell-culture supernatants were collected, and fresh media containing M-CSF and/or ATRA was added every 3 days. **(D-G)** MDMs exposed to HIV_NL4.3BaL_ (**D-E**) and T/F HIV_THRO_ (**F-G**) were analysed for HIV-DNA integration by nested real-time PCR at day 3 post-infection (**D and F**) and viral replication by HIV-p24 ELISA every 3 days up to 15 days post-infection (**E and G**). Shown are the kinetics of HIV-1 replication in one representative donor (**E and G, left panels**) and statistical analysis performed at day 9 post-infection for n=8 participants (**E and G, middle and right panels)**. Wilcoxon p-values are indicated on the graphs.

For experiments depicted in Figure 1, the optimal concentration of ATRA (10 nM) was determined in preliminary dose-response studies on MDMs infected with HIV_NL4.3BaL_. This concentration proved to consistently increase HIV-1 replication in MDMs in all donors tested (Supplemental Figure 2A-B), with no influence on cell viability, as observed by flow cytometry (Supplemental Figure 2C). This concentration is also within the range of physiological plasma RA levels (4-14 nM)^79-81^ and was previously demonstrated by our group to efficiently boost HIV-1 replication in CD4^+^ T-cells^60,72-74^.

To explore whether ATRA preferentially affects R5 HIV-1 replication, MDMs were exposed to the replication-competent CXCR4-tropic (X4) HIV_NDK_. In contrast to R5 HIV-1, ATRA did not induce significant changes in HIV-1 integration and replication upon exposure to ATRA (Supplemental Figure 3A-B).

Together, these results demonstrate that ATRA render MDMs highly permissive to productive R5 but not X4 HIV-1 infection, a process likely explained by the differential modulation of CCR5 and CXCR4 by ATRA, leading to a more efficient CCR5-mediated viral entry.

### ATRA increases HIV-1 permissiveness in MDMs at post-entry and post-integration levels

To identify the post-entry steps of the viral replication cycle modulated by ATRA, MDMs were exposed to single-round VSV-G-pseudotyped HIV-1 (HIV_VSV-G_) expressing *gfp* in place of *nef* ^73^, a viral construct capable of entering cells independently of CD4/CCR5/CXCR4, *via* the low-density lipoprotein receptor (LDLR)^82^. In a first set of experiments, MDMs were treated with ATRA before HIV_VSV-G_ exposure (Figure 2A). At day 3 post-infection, levels of early (RU5) and late (Gag) reverse transcripts, as well as levels of integrated HIV-DNA (proviral HIV-DNA), measured as we previously reported^73,76,83^, were significantly increased by ATRA (Figure 2B), indicative of a superior efficacy in reverse transcription and/or integration. Consistently, HIV-p24 levels in cell-culture supernatants, indicative of viral production, were significantly increased by ATRA (n=14; p=0.0009) (Figure 2C). These results indicate that ATRA acts post-entry on early steps of HIV replication cycle between reverse transcription and integration leading to efficient subsequent viral production.

**Figure 2:**
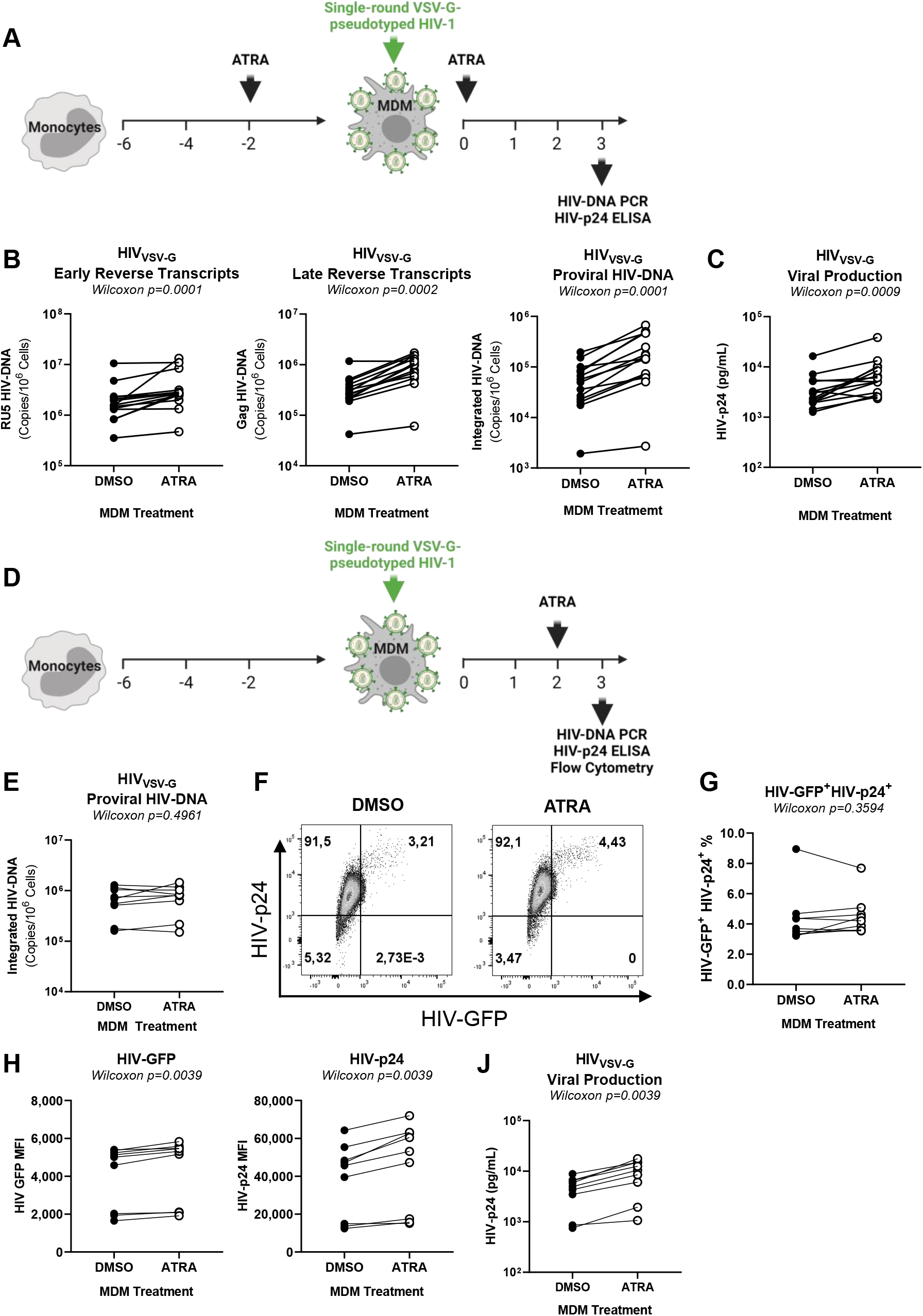
ATRA increases HIV-1 replication at post-entry levels, before and after integration. **(A)** Shown is the experimental flowchart, when MDMs were exposed to single-round VSV-G pseudotyped HIV-1 (HIV_VSV-G_) and treated with ATRA (10nM) before and after infection **(A-C)**. Briefly, MDMs and cell-culture supernatants were harvested at day 3 post-infection for HIV-DNA and HIV-p24 quantification, respectively. Shown are the levels of early reverse transcripts (RU5) **(B, left panel)**, late reverse transcripts (Gag) **(B, middle panel),** and integrated HIV-DNA (Alu/LTR) **(B, right panel)**, as well as HIV-p24 levels in cell culture supernatants **(C)**. Shown is the experimental flowchart when MDMs were exposed to ATRA (10nM) at day 2 post-infection **(D)**. At day 3 post-infection, cells were collected and analyzed for HIV-DNA integration by real-time nested PCR and for GFP and intracellular HIV-p24 expression by flow cytometry. Shown are statistical analysis of HIV-DNA integration **(E)**, GFP and HIV-p24 co-expression in MDMs from one representative donor **(F)**, as well as statistical analysis of the frequency of infected (GFP^+^HIV-p24^+^) MDMs **(G)**, and the GFP **(H, left panel)** and HIV-p24 **(H, right panel)** MFI expression from n=9 participants. Finally, shown are statistical analysis of HIV-p24 levels in cell-culture supernatants measured by ELISA **(J)**. Experiments were performed on MDMs from n=14 **(A-C)** and n=9 (**D-J**) HIV-uninfected participants. Wilcoxon values are indicated on the graphs.

Considering the fact that RARE are present in the HIV-LTR and promote HIV transcription^69^, we aimed to study the effects of ATRA in MDMs at post-integration level. To this aim, in a second set of experiments, MDMs were treated with ATRA at day 2 post-infection (Figure 2D), a time when HIV integration is maximal^84^. As expected under these conditions, there were no differences in HIV-DNA integration (n=9; p=0.4961) (Figure 2E) nor in the frequency of productively infected GFP^+^HIV-p24^+^ cells (n=9; p=0.3594) (Figure 2F-G) in ATRA-MDMs *versus* DMSO-MDMs. However, exposure to ATRA post-integration significantly increased the MFI for both GFP and HIV-p24 expression (n=9 p=0.0039) (Figure 2H), as well as viral release (n=9; p=0.0039) (Figure 2J), indicative of an efficient transcription/translation in the presence of ATRA.

Overall, these results demonstrate that ATRA render MDMs highly permissive to HIV-1 replication at various post-entry levels, between reverse transcription and integration, as well as post-integration, likely by promoting HIV transcription and/or virion production in MDMs.

### ATRA transcriptionally reprograms MDMs for increased permissiveness to HIV replication

To gain molecular insights into mechanisms underlying the effects of ATRA on HIV-1 replication in MDMs, genome-wide RNA sequencing was performed prior HIV-1 exposure (Figure 3A). Differentially expressed genes (DEG) were identified based on p-values (<0.05), adjusted p-values (<0.05), and fold change (FC, cut-off of 1.3) (Supplemental Figure 4A), with 1,772 and 2,047 transcripts identified as being upregulated and downregulated, respectively, in ATRA-MDMs *versus* DMSO-MDMs (<u>Supplemental Files 1-2</u>).

**Figure 3:**
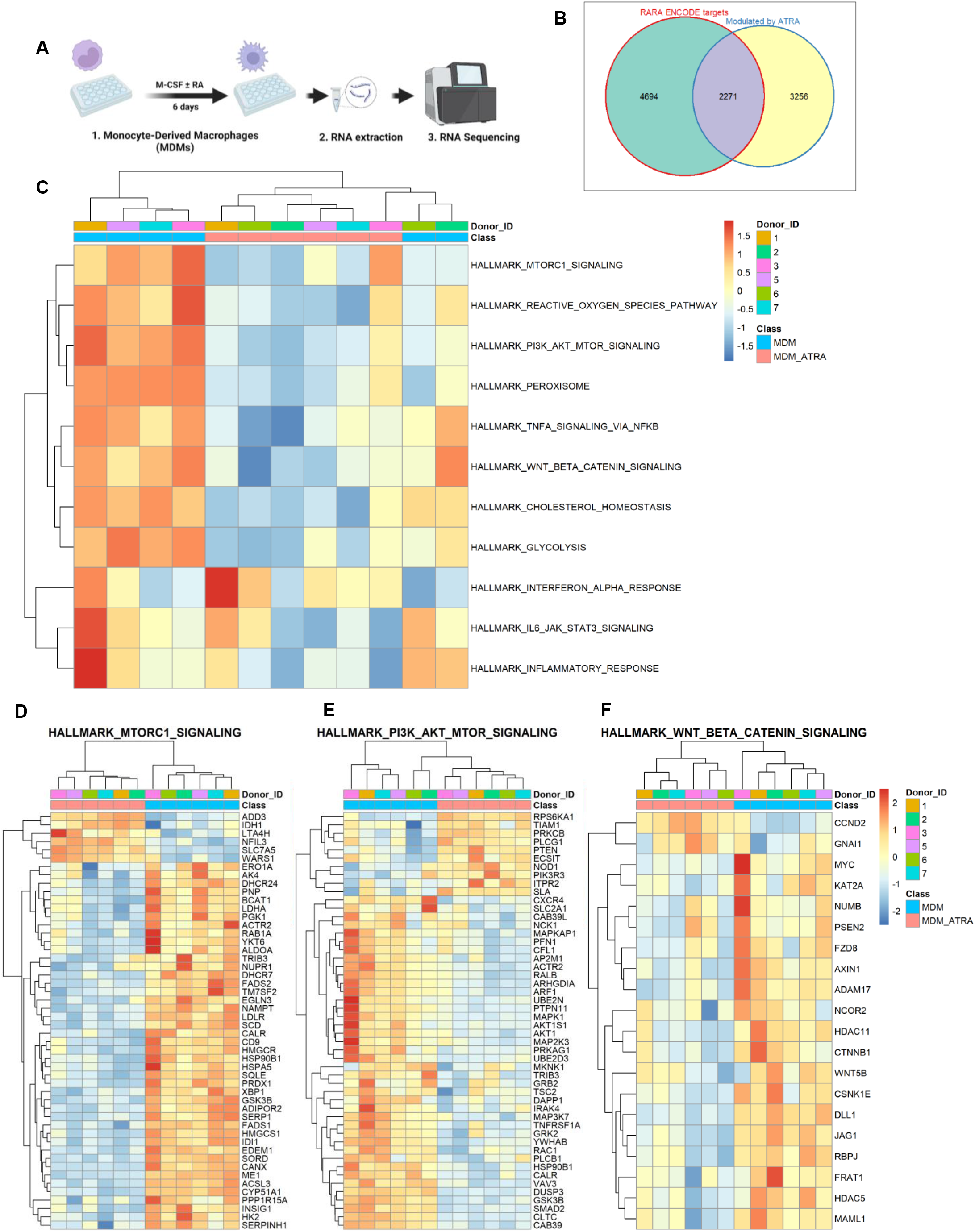
ATRA transcriptionally reprograms MDMs and specifically modulates mTOR and Wnt/β-Catenin signaling pathways. **(A)** Shown is the experimental flowchart. Total RNA extracted from MDMs of six HIV-uninfected participants generated in the presence (ATRA- MDMs) or the absence of ATRA (10nM) (DMSO-MDMs) were used for RNA sequencing. **(B)** Differentially expressed genes (DEG) were analyzed for the presence of RA-responsive elements (RARE) in their promoters, using the ENCODE Bioinformatic tool (https://www.encodeproject.org). This allowed the identification of n=2,271 DEGs that may represent putative direct RA transcriptional targets in ATRA-treated MDMs. **(C)** Further, Gene set variation analysis (GSVA) was performed to identify signaling pathways modulated by ATRA in MDMs. Heatmaps depict top modulated signaling pathways in ATRA-treated MDMs **(C**), as well as gene sets associated with three top modulated pathways: mTORC1 **(D)**, PI3K AKT mTOR **(E)**, and Wnt/β-Catenin **(F).**

Among the top 50 modulated genes, *HIVEP2* (an enhancer of HIV transcription^85^), *IRF1* (a facilitator of HIV replication^86^), *ABI3* (an inducer of cell senescence^87^), *TTC7A* (a modulator of chromatin structure and regulator of HIV transcription^88^), *PHOSPHO1* (a regulator of energy metabolism^89^), and *RUNX2* (a modulator of macrophage differentiation^90^ and a negative regulator CXCR4 expression^91^) were upregulated, while *HIVEP1* (a negative regulator of NF-κB activation^92^), *DUSP3* (a mediator of LPS and macrophage polarization^93^), and *SLC11A1* (encoding for natural resistance-associated macrophage protein 1, with polymorphism associated with HIV mortality^94^) were downregulated in ATRA-MDMs *versus* DMSO-MDMs (Supplemental Figure 4B). These transcriptional changes may explain the increased permissiveness to productive R5 HIV-1 replication in ATRA-MDMs, with decreased CXCR4 expression and increased viral transcription.

#### Identification of direct RA target genes

Among DEG modulated by ATRA in MDMs, 2,271 transcripts (*e.g., CEACAM1, ITGA1, BATF2, HIVEP2, ITGAE, WARS1, IRF1, CD7, GFI1, NFIL3, ITGA6*, *CD55*), expressed RA responsive elements (RARE) in their promoters, as identified using the *in silico* Bioinformatics tool Encode search (https://www.encodeproject.org) (Figure 3B; <u>Supplemental File 3</u>), indicative that ATRA directly regulates the expression of specific genes in MDMs.

#### Gene Set Variation Analysis (GSVA)

To extract further meaning from these transcriptional changes, GSVA (C2, C3, C5, C7, C8, and Hallmark databases; https://www.gsea-msigdb.org/gsea/msigdb/index.jsp) was performed. Top-modulated hallmark pathways are depicted in Figure 3C. Components and regulators of the mTORC1 signaling, PI3K/Akt/mTOR signaling, and Wnt/β-catenin signaling are illustrated in Figures 3D-F and detailed here below:

#### mTORC1 Signaling

Among transcripts linked to the mTOR pathway, the expression of amino acid transporters such as *SLC7A5*^95^ and Tryptophanyl-tRNA synthetases 1 such as *WARS1*^96^ are upregulated suggesting an increase in protein synthesis and metabolic activity (Figure 3D). Moreover, the expression of certain negative regulators of mTOR, such as *BCAT1^97^*, *TRIB3*^98^, *NUPR1*^99^, *NAMPT*^100^, and *PRDX1*^101^ was downregulated by ATRA in MDMs, indicative of mTOR activation (Figure 3D). Furthermore, the expression of the insulin-induced gene 1 (*INSIG1)*, a regulator of lipid metabolism^102^ and an HIV-1 restriction factor involved in Gag degradation^103^, was downregulated by ATRA in MDMs (Figure 3D). Overall, these results are indicative of an mTOR-dependent ATRA-mediated reprogramming of MDMs metabolic activity for increased HIV-1 permissiveness.

#### PIK3-AKT-mTOR Signaling

The activation of mTOR pathway requires the activation of several kinases such as phosphoinositide 3 kinase (PIK3) and protein kinase B (AKT)^104^. The class IA PI3K consists of heterodimers of the p110 catalytic (PIK3CA, PI3KCB or PI3KCD) subunit and p85 regulatory subunit (PIK3R1, PIK3R2, and PIK3R3)^105^. Results in Figure 3E demonstrate that the expression of the p85 regulatory subunits (*PI3KR3*) was upregulated, while the expression of *AKT,* encoding for a kinase upstream of mTOR^104^. was downregulated by ATRA in MDMs. The latter coincided with the increased expression of the *phosphatase and tensin homolog* (*PTEN*) (Figure 3E), a negative regulator of PI3K and AKT^104^. The expression of *tuberous sclerosis 2* (*TSC2*), a negative regulator of mTOR ^104^, was downregulated, while the expression of kinases downstream of mTORC1, such as the ribosomal protein S6 kinase 1 (*RPS6KA1*)^104^, was upregulated (Figure 3E), suggesting the activation of mTORC1 pathway in ATRA-MDMs *versus* DMSO-MDMs. Importantly, S6K inhibits the PI3K-AKT-mTOR pathway through a negative feedback loop^104^, which is consistent with the higher expression of *PTEN* and lower expression of *AKT* genes in ATRA-MDMs *versus* DMSO-MDMs (Figure 3E). Moreover, the expression of the protein kinase C (*PRKCB*), another kinase downstream mTORC2^104^, was upregulated in ATRA-MDMs *versus* DMSO-MDMs (Figure 3E), suggesting the activation of mTORC2 through the activation of AKT^104^, Furthermore, the expression of *CAB39*, responsible for activating AMPK, a kinase that inhibits mTOR activation through sensing of cellular energy (AMP:ATP ratio^104,106^, was also downregulated in ATRA-MDMs compared to DMSO-MDMs (Figure 3E). Overall, these results are indicative of ATRA activating the PI3K-AKT-mTOR pathway in MDMs, thus facilitating HIV-1 replication.

#### Wnt/β-catenin

The Wnt/β-catenin pathway controls key cellular functions, such as cell proliferation, differentiation, and apoptosis, and negatively regulates HIV-1 transcription^107^. Results in Figure 3F depict the downregulation of *FZD8 (Wnt receptor)*, *AXIN1 (a component of the β- catenin/APC/GSK3β complex)*, *CTNNB1* (β-catenin), and *WNT.5B* transcripts. Other components/regulators of the Wnt/β-catenin pathway are listed among DEGs (<u>Supplemental</u> <u>Files 1-2</u>). Of particular notice, components of the canonical Wnt/β-catenin pathway are downregulated (*i.e., MYC, PDK1, MMP*) <u>(Supplemental File 2</u>), suggesting the inactivation of this pathway ^108^. Furthermore, transcripts linked to the non-canonical Wnt/β-catenin pathway were upregulated (*i.e., PLC, PRKCB, NFATC2*) in ATRA-MDMs *versus* DMSO-MDMs (<u>Figure 3E and Supplemental File 1</u>)^109^, pointing to the activation of the non-canonical Wnt/β-catenin pathway. The expression of *TCF4/TCF7L2*, a gene encoding for the transcription factor TCF4 that inhibits HIV transcription through its binding on the 5’HIV LTR promoter upon association with β-catenin^110^, was upregulated (<u>Supplemental File 1</u>) and transcripts for *GSK3B*, a serine-threonine kinase that mediates the proteolytic degradation of β-catenin^108^, were downregulated by ATRA in MDMs (Figure 3D-E). While the ATRA-mediated upregulation of *TCF4/TCF7L2* and the downregulation of *GSK3B* transcripts remains intriguing, the downregulation of β- catenin expression is consistent with high levels of HIV-1 replication in ATRA-MDMs.

These results point to profound transcriptional reprogramming mediated by ATRA in MDMs and reveal two druggable pathways, mTOR and Wnt/beta-catenin, that may play a key role in this process.

### ATRA modulates the expression of HIV interactors in MDMs

To extract further meaning from the large sets of RNA-Sequencing data, the NCBI HIV interactor database was interrogated for known HIV permissiveness/restriction factors modulated by ATRA in MDMs (Figure 4A-B). First, ATRA increased *CCR5* and decreased *CD4* mRNA expression in MDMs (Figure 4A), consistent with the flow cytometry results in Figure 1B-C. Among transcripts known to increase HIV replication, ATRA increased the expression of the transcription factor *NFATC2*^111^ and coactivator *NCOA3*^112^ (Figure 4A-B). At the opposite, ATRA decreased expression of *BIRC2* (suppressor of HIV transcription)^113^, *SERINC5* (restriction factor for HIV-1 release)^114^, *RNASE1* (inhibitor of HIV-1 production)^115^, *SOD2* (an integral component of the cellular antioxidant system)^116^, *HDAC7* (suppressor of HIV transcription)^117^, and *GSK3B (an inducer of zink-finger antiviral protein (ZAP) functions*)^118^ (Figure 4A-B). Unexpectedly, genes encoding for positive regulators of HIV transcription (*TRIM32*^119^, *NCOA1*^120^) and virion production/release (*LGALS3*^121^, *FURIN*^122^. Furthermore, *HIC1* and *TCF4/TCF7L2*^123^ (Figure 4A-B), two negative regulators of HIV transcription^124^, were upregulated by ATRA in MDMs. RT-PCR validations confirmed the increased expression of *HIC1* mRNA in ATRA-exposed MDMs (Supplemental Figure 5A). At the opposite, the expression of *PPARG,* a HIV-1 transcriptional repressor^125^, was downregulated in MDMs upon exposure to ATRA (<u>Supplemental File 2</u>), as validated by RT-PCR quantification (Supplemental Figure 5B).

**Figure 4:**
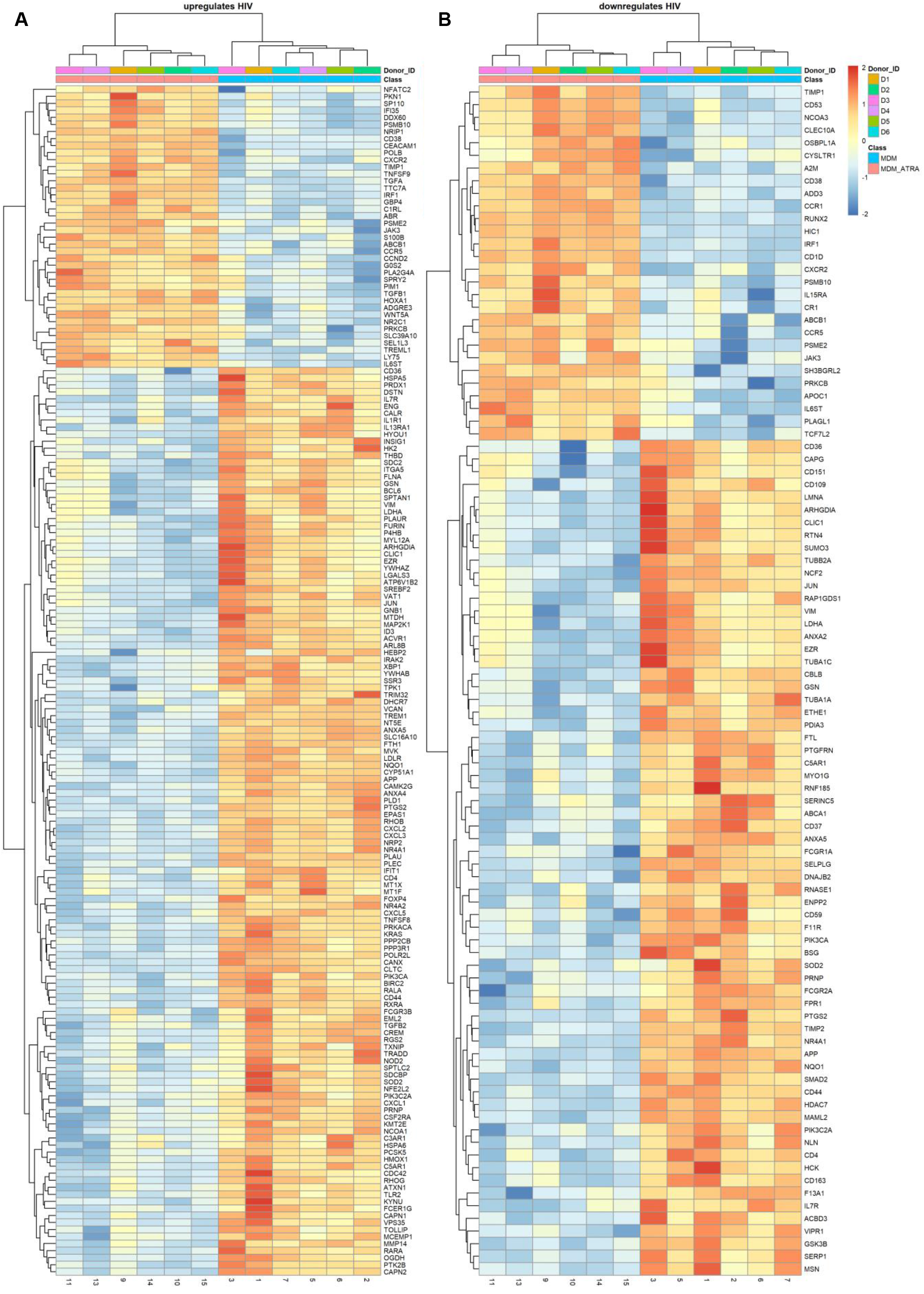
Meta-analysis of genes modulated by ATRA in MDMs included in the NCBI HIV interaction database. DEGs were identified as in <u>Figure 3</u>. Shown are transcripts modulated by ATRA in MDMs (p<0.05; FC cut-off 1.3) matching the lists of human genes included on the NCBI HIV interaction database involved in HIV upregulation **(A)** and downregulation **(B)**. Heatmap cells are scaled by the expression level z-scores for each probe individually. Results from each donor are indicated with a different color code (n=6).

Together, these results reveal a panel of HIV-1 restriction and dependency factors, with a fraction of them identified as genes with putative RARE in their promoters (*e.g., SAMHD1, CTNB1, GSK3B,* SLC7A5, *WARS1, TRIB3, NUPR1, NAMPT, PRDX1, AKT1, PTEN, HIVEP2, HIVEP1, SERINC5*) (<u>Supplemental File 3</u>). Some of these transcripts are linked to mTOR and Wnt/β-catenin pathways, thus providing a molecular explanation for the dramatic increase in HIV-1 replication in ATRA-treated MDMs at entry and post-entry levels. Other genes upregulated by ATRA (Supplemental Figure 1) expressing RARE in their promoters and reported to contribute to HIV-1 replication were identified using another *in silico* search method on www.genecards.org and included CCR5, MTOR, RPS6KB1, TCF7L2, and INSIG1.

### ATRA increases HIV-1 permissiveness in MDMs *via* mTOR-dependent mechanisms

To explore the role of the mTOR pathway in modulating HIV-1 permissiveness in ATRA-treated MDMs (Figure 3D-E), the expression of total and phosphorylated mTOR was visualized by western blotting. Results in Supplemental Figure 6A-B demonstrated an increased expression of total and phosphorylated mTOR in ATRA-treated MDMs from n=3 HIV-uninfected donors. Consistently, the expression of total and phosphorylated S6K, a kinase downstream mTOR indicative of mTOR activity^104^, was also increased in ATRA-MDMs *versus* DMSO-MDMs (Supplemental Figure 6C-D). This prompted us to explore the contribution of mTOR activation on HIV-1 permissiveness in ATRA-treated MDMs. To this aim, MDMs generated in the presence/absence of ATRA were first analyzed for the expression of CD4, CCR5, and CXCR4 upon exposure to INK128, a documented mTOR inhibitor^126^. The expression of CD4 and CCR5 was significantly reduced on both DMSO-MDMs and ATRA-MDMs, while the CXCR4 expression was increased on DMSO-MDMs upon exposure to INK128 (Figure 5A-B). Similarly, exposure to INK128 significantly decreased HIV_THRO_ integration and replication, mainly in ATRA-MDMs (Figure 5C-D). Consistently, exposure to INK128 resulted in a significant decrease in the expression of total and phosphorylated mTOR, as well as total and phosphorylated S6K in ATRA-treated MDMs, as visualized by western blotting (Supplemental Figure 7A-D). These results demonstrate the capacity of INK128 to act on the mTOR activation pathway in ATRA-MDMs and limit R5 HIV-1 replication, in part, by decreasing CD4 and CCR5 but not CXCR4 expression.

**Figure 5:**
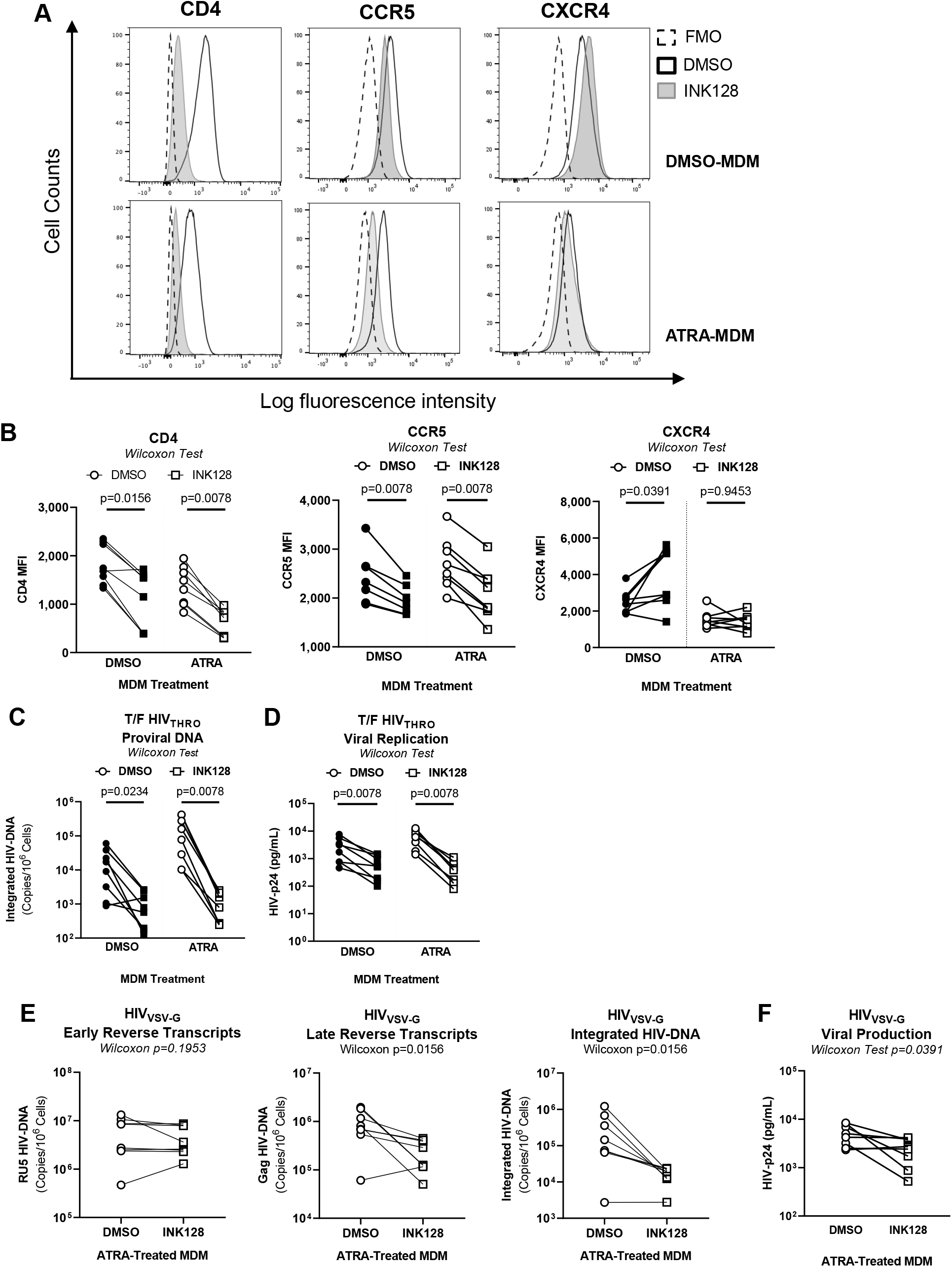
mTOR inhibition counteracts the effect of ATRA on CCR5 expression and HIV replication and integration in MDMs. MDMs were generated in the presence/absence of ATRA (10 nM), as in Figure 1A, and treated with the mTOR inhibitor INK128 (50 nM) two days before infection and the day of infection. **(A)** Shown are the representative flow cytometry histograms of extracellular CD4, CCR5 and CXCR4 expression. **(B)** Shown are statistical analysis of relative MFI of CD4 **(left panel)**, CCR5 **(middle panel)**, and CXCR4 **(right panel)** expression. **(C-D)** A fraction of ATRA-MDMs and DMSO-MDMs pretreated or not with INK128 were exposed to replication-competent CCR5-tropic T/F HIV_THRO_. Cells and cell-culture supernatants were harvested on day three post-infection for the quantification of integrated HIV-DNA by nested real-time PCR **(C)** and soluble HIV-p24 by ELISA **(D)**. **(E-F)** In parallel, another fraction of ATRA-MDMs and DMSO-MDMs pretreated with INK128 were exposed to a single-round VSV-G-pseudotyped HIV constructs (HIV_VSV-G_) for three days. Shown are the levels of early reverse transcripts **(RU5; E, left panel)**, late reverse transcripts **(Gag; E, center panel)**, integrated HIV-DNA **(Alu; E, right panel)**, as well as the levels of HIV-p24 **(F)**. Experiments were performed on MDMs from n=8 HIV-uninfected individuals. Wilcoxon values are indicated on the graphs.

To identify the viral replication steps targeted by mTOR in ATRA-MDMs, a single-round infection was performed with HIV_VSV-G_. It is noteworthy that INK128 significantly decreased late reverse transcripts (Gag) and proviral HIV-DNA, but not early reverse transcripts (RU5), indicative of mTOR-dependent mechanisms affecting the completion of reverse transcription and subsequently integration, but not the initiation of reverse transcription (Figure 5E). Consistently, results in Figure 5F demonstrate that INK128 also reduces virion release in ATRA-treated MDMs in this single-round model of infection. Together, these results demonstrate that INK128 can counteract the effects of ATRA on CCR5 expression, HIV reverse transcription and integration, and virion release, indicative that ATRA promotes HIV-1 permissiveness in MDMs *via* mTOR-dependent mechanisms.

### ATRA modulates SAMHD1 phosphorylation *via* mTOR-dependent mechanisms

The mTOR interferes with HIV-1 replication at multiple steps of the viral replication cycle, including reverse transcription^127,128^. One restriction factor originally identified for its capacity to restrict HIV-1 replication in myeloid cells by interfering with the completion of reverse transcription is SAMHD1^129^. Interestingly, SAMHD1 was found upregulated by ATRA and present RARE in its promoter (<u>Supplemental Files 1 and 3</u>). The potential link between mTOR and SAMHD1 activity is also supported by the upregulated expression of CCND2 (<u>Supplemental</u> <u>File 1,</u> Figure 6A) encoding for cyclin D2, a modulator of cell cycle and SAMHD1 activity^130,131^. Indeed, phosphorylation via cyclin dependent kinases (CDK) 1/2 was reported to reduce SAMHD1 capacity to restrict HIV in macrophages^131^. In line with this prediction, results in Figure 6B-D demonstrate an increased expression of total and phosphorylated SAMHD1 in MDMs upon exposure to ATRA. Oppositely, exposure to INK128 led to a significant decrease in SAMHD1 phosphorylation mainly in ATRA-MDMs (Figure 6B-D), in line with the antiviral activity of INK128 (Figure 5C-F). These results demonstrated the regulation of SAMHD1- mediated HIV restriction in MDMs *via* mTOR-dependent mechanisms.

**Figure 6.**
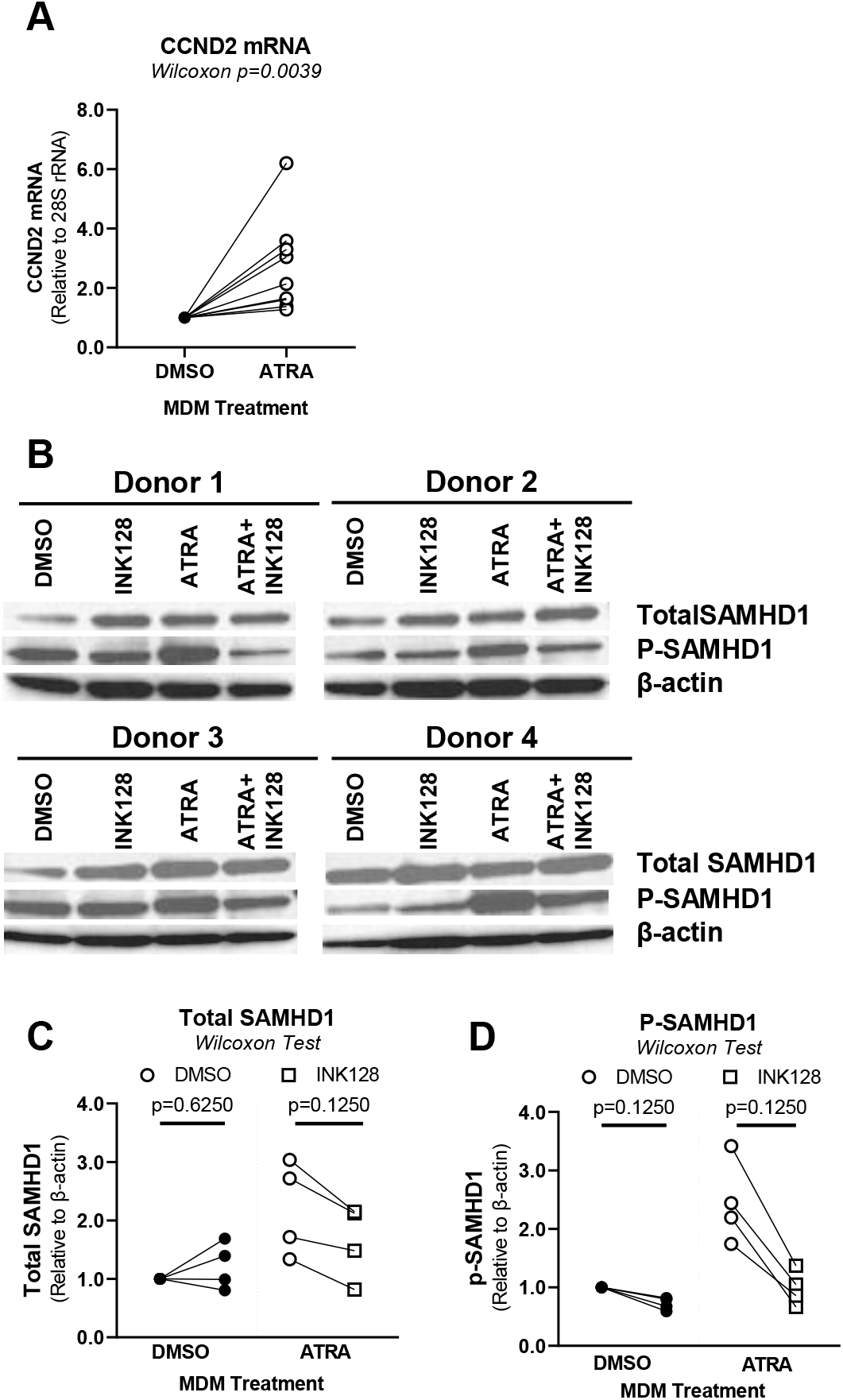
ATRA modulates the expression of Cyclin D2 and SAMHD1 phosphorylation in a mTOR-dependent manner. **(A)** MDMs were generated in the presence/absence of ATRA (10nM) and pre-treated or not with INK128 (50 nM), as in Figure 5. Prior HIV-1 infection, cells were harvested for RT-PCR **(A)** and western blotting investigations **(B-D)**. Shown are levels of Cyclin D2 (CCND2) mRNA expression (**A)**, as well as levels of total (MW: 72 KDa) and phosphorylated (MW: 72 KDa) SAMHD1, and β-actin (MW: 42 KDa) expression **(B)** in MDMs from four different HIV-uninfected donors. **(C-D)** Graphs depict total (**C)** and phosphorylated SAMHD1 levels **(D)** normalized to β-actin levels. Experiments were performed on MDMs from n=9 HIV-uninfected individuals. Wilcoxon values are indicated on the graphs.

### The **β**-catenin/TCF4 pathway limits excessive HIV-1 replication in ATRA-MDMs

Our RNA-Sequencing revealed a decreased expression for *CTNNB1* (encoding for β-catenin) and *GSK3B*, and an increased expression of *TCF7L2* (encoding for TCF7L2/TCF4) in ATRA- MDMs *versus* DMSO-MDMs (<u>Supplemental Files 1-2</u>). Here, we first validated the upregulation of TCF4 mRNA and protein expression in ATRA-treated MDMs (Supplemental Figure 8A-C), as well as the downregulation of CTNNB1/β-catenin (Figure 7A-C) and GSK3β expression (Figure 7D-F) at mRNA and protein level in ATRA-MDMs *versus* DMSO-MDMs. Then, to explore the role of β-catenin/TCF4 in regulating HIV_THRO_ replication in ATRA-treated MDMs, ATRA-MDMs were exposed to PRI-724, a potent inhibitor that disrupts the interaction between β-catenin and CBP^132^, and PNU-74654, a potent inhibitor that disrupts the interaction between β- catenin and TCF4^133^. In preliminary experiments, the optimal concentrations of PNU-724 (0.1 μM) or PNU-74654 (5 μM) were selected based on dose-response experiments measuring cell viability and HIV-1 replication (<u>data not shown</u>). Levels of HIV-DNA integration were further significantly increased in ATRA-MDMs when HIV_THRO_ infection was performed in the presence of PRI-724 (Figure 7G) or PNU-74654 (Figure 7H). Together, point to β-catenin/TCF4 as important negative regulators of HIV-1 replication in ATRA-treated MDMs. This negative feedback antiviral mechanism, likely limits exacerbated HIV-1 replication in macrophages in an environment rich in RA.

**Figure 7:**
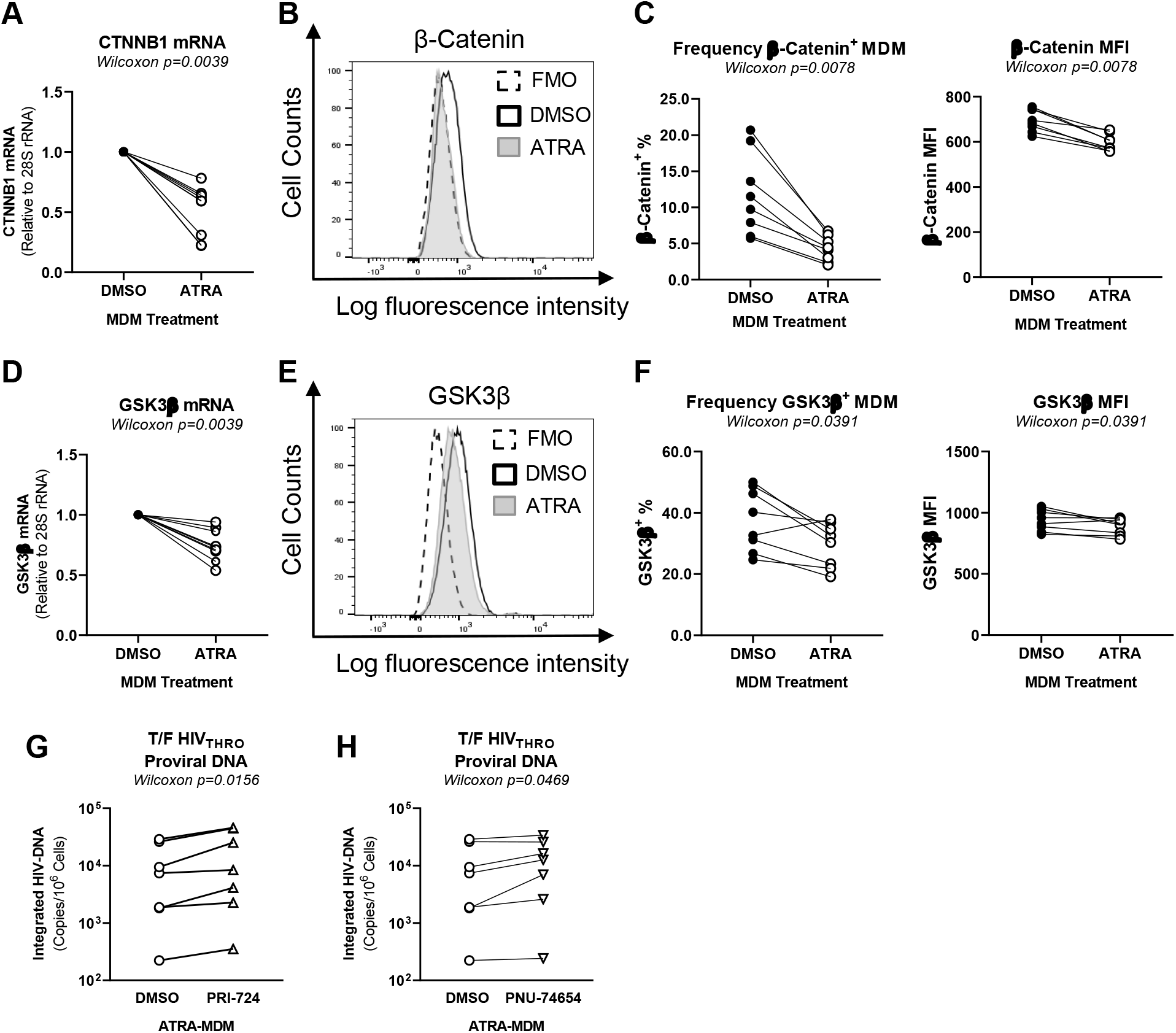
The Wnt/β-catenin pathways blockade further increase HIV-1 replication in ATRA-treated MDMs. MDMs were generated in the presence/absence of ATRA (10 nM) as in Figure 1A and exposed to PRI-724 (0.1 µM) or PNU-74654 (10 µM) two days before infection and the day of infection. MDMs harvested prior HIV-1 infection were used for the analysis of β- Catenin **(A-C)** and GSK3β **(D-F)** mRNA expression by RT-PCR (**A** and **D**) and intranuclear staining using specific Abs and flow cytometry analysis (**B-C** and **E-F**). Shown is the statistical analysis of β-Catenin **(A)** and GSK3β **(D)** mRNA expression in MDMs (n=8). Shown are flow cytometry histograms for β-Catenin and GSK3β expression on MDMs from one representative donor **(B, E)**, as well as statistical analysis of the β-Catenin **(C)** and GSK3β **(F)** expression, in terms of % **(left panels)** and MFI **(right panels)** in MDMs (n=8). **(G-H)** Finally, ATRA-MDMs were pretreated with the β-catenin inhibitors PRI-724 and PNU-74654 and then exposed to T/F HIV_THRO._ At day 3 post-infection, levels of integrated HIV-DNA were quantified by nested real-time PCR in ATRA-MDMs pre-treated with PRI-724 **(G)** and PNU-74654 **(H)**. Shown are statistical analysis performed with results generated with MDMs from n=7 donors. Wilcoxon values are indicated on the graphs.

## DISCUSSION

Macrophages are major players in host-pathogen interactions, with a well-established division of labor between long-lived embryonically derived self-renewing TRMs, mainly involved in tissues remodeling and homeostasis, and short-lived MDMs, which contribute to inflammatory processes^50^. Most recent advances support a scenario in which HIV-1 exploits this functional dichotomy for its dissemination before ART initiation and for VR persistence *via* direct/indirect mechanisms in ART-treated PLWH^12,33-41^. Of particular notice, the ageing process is associated with the progressive replacement of TRMs by MDMs, as originally demonstrated for heart macrophages^53,54^. Therefore, studies on factors regulating HIV-1 replication in MDMs become highly relevant in the context of premature ageing reported in ART-treated PLWH ^134,135^.

In this manuscript, we report that RA render MDMs highly permissive to HIV-1 replication *via* multiple entry/post-entry mechanisms governed by mTOR, as demonstrated by the use of the mTOR inhibitor INK128. mTOR is a serine/threonine protein kinase that regulates cell metabolism in response to changes in the environment, such as nutrients and growth hormones^128^. In line with the current findings, we previously reported that ATRA acts on Th17- polarized CCR6^+^CD4^+^ T-cells to increase their permissiveness to HIV-1 infection *in vitro* and boosts viral outgrowth from cells of ART-treated PLWH *via* mTOR-dependent mechanisms^60,72-74^. Other groups previously demonstrated the key role played by mTOR in the positive regulation of HIV-1 replication in CD4^+^ T-cells^128^. Taylor *et al.,* demonstrated that increased glycolysis, through the modulation of mTOR, can facilitate HIV-1 permissiveness through increasing dNTP pools required for early steps of HIV-1 replication as well as acetyl-CoA required for nuclear transportation of viral products^127^. Ono *et al*., have demonstrated that increased lipid synthesis, through the modulation of mTOR, can complete the viral life cycle by increasing the level of cholesterol and membrane lipids required for the late steps of HIV-1 budding^136^. Besnard *et al.* demonstrated that both mTORC1 and mTORC2 are essential for HIV-1 reactivation from latency^137^ and suggested that mTOR inhibition activates the autophagy pathway, which has been demonstrated to repress HIV production through Tat degradation in CD4^+^ T cells^138^. Also, Heredia *et al.* have reported that INK128-mediated mTOR inhibition decreased CCR5 expression and HIV transcription^139^. This evidence supports the link between viral replication and host cell metabolism^140^. In this context, mTOR inhibitors, used in clinic for the treatment of diabetes and tested in clinical trials for cancer^141,142^, may also be used in HIV-1 infection. Most recently, our group and others provided evidences pointing to metformin, an indirect mTOR inhibitor, as an efficient therapeutic strategy for the control of residual HIV-1 transcription and premature ageing in ART-treated PLWH^128,143,144^.

Of particular notice, ATRA upregulated CCR5 but downregulated CD4 and CXCR4, and preferentially promoted R5 HIV replication in MDMs. This contrasts with results on CD4^+^ T- cells where ATRA similarly boosted R5 and X4 HIV-1 strain replication^73^. These results raise new questions on the role of ATRA in the selection of R5 HIV-1 strains during primary infection in MDMs, a mechanism essential for the establishment of transmitted founder HIV-1 reservoirs^145^. The fact that ATRA decreased CD4 expression in MDMs without impeding on the efficacy of productive HIV-1 infection is consistent with the reported capacity of HIV-1 to enter macrophages with low CD4 requirement^146^. The upregulation of CCR5 in ATRA-treated MDMs was in line with our *in silico* findings upon ENCODE database interrogation that RARE are present in the CCR5 promoter, suggesting that CCR5 is a direct transcriptional target for ATRA. Molecular mechanisms by which the mTOR inhibitor INK128 counteracted the effect of ATRA on CCR5 expression remain to be elucidated.

Single-round infection with VSV-G-pseudotyped HIV-1 demonstrated that ATRA acts also at post-entry levels, facilitating the efficacy of reverse transcription and leading to an efficient HIV-DNA integration. Genome-wide RNA sequencing, GSVA, NCBI HIV interactor data base search, and the in silico ENCODE search for genes with RARE in their promoters, together with functional validations and pharmacological targeting allowed us to identify ATRA as an inducer of SAMHD1 phosphorylation, a post-translational modification associated with the loss of its capacity to restrict HIV-1 reverse transcription ^147^. SAMHD1 was originally identified as a key HIV-1 restriction factor in macrophages; its mechanism of action involved the control of the pool of dNTP essential for reverse transcription^148,149^. In the current studies, ATRA-induced SAMHD1 phosphorylation coincided with increased expression of the cyclin CCND2. Other studies reported the role of CDK1/2 in the phosphorylation of SAMHD1^131^. We here demonstrate that ATRA-mediated SAMHD1 phosphorylation was abrogated by the mTOR inhibitor INK128. This knowledge adds to the beneficial effects of mTOR inhibitors on controlling HIV-1 infection, mainly in the context were the antiviral features of SAMHD1 are linked to VR and disease progression in PLWH^148,149^.

In the context of single-round infection with VSV-G-pseudotyped HIV-1, MDM exposure to ATRA upon optimal HIV-DNA integration demonstrated an increased intracellular expression of HIV-p24 and an increased virion release in cell culture supernatants. This points to an effect of ATRA on HIV-1 transcription and/or translation. The effect of ATRA on HIV-1 transcription can be explained by the presence of RARE in the HIV-LTR^69^. Moreover, ATRA activated mTOR/S6K *via* phosphorylation, a pathway documented to promote HIV-1 transcription *via* the modulation of CDK9/p-TEFb complex, a component of the viral transcriptional machinery^137,150^. Furthermore, our RNA sequencing results originally revealed that ATRA decreased the expression of β-catenin and GSK3β, while increasing the expression of TCF7L2/TCF4 in MDMs. Wnt/β-catenin was recognized as a restriction factor of HIV^151^. The Wnt/β-catenin pathway represses HIV-1 transcription *via* mechanisms involving the β-catenin interaction with TCF4 and subsequent binding on HIV-LTR^107,110,123^ or the suppression of transcription factors that increase HIV transcription, namely C/EBP and NF-κβ^151^. In the absence of Wnt signaling, GSK3β promotes the proteasomal degradation of β-catenin^108^. GSK3β was also reported to facilitate TAT-mediated HIV-1 transcription. Therefore, by decreasing the expression of β- catenin and GSK3β at mRNA and protein levels, ATRA may facilitate HIV-1 transcription. Nevertheless, the Wnt/β-catenin/TCF4 pathway remained functional in ATRA-treated MDM, likely as a mechanism to limit exacerbated HIV-1 replication, as demonstrated by the increased expression of TCF7L2/TCF4 mRNA/protein and the use of the β-catenin inhibitors PRI-742 and PNU-74654. In line with this idea, CD4^+^ T cells from elite controllers exhibit higher expression/activation of the Wnt/β-catenin pathway compared to ART-treated patients, explaining limited viral rebound^152^. Other studies documented the role of Wnt/β-catenin in CD4^+^ T cells protection from HIV-mediated apoptosis^153^ and demonstrated that PRI-724 limited T cell proliferation *in vivo,* without reducing the VR size^154^. Finally, Barbian *et al*., have demonstrated that inhibiting the Wnt/β-catenin pathway reactivates HIV in primary CD4^+^ T cells from ART- treated PLWH and increases the activity of latency-reversing agents^107^. The interplay between the mTOR and Wnt/β-catenin pathways was documented^155^, with mTOR regulating the expressionWnt receptor, FZD^156^, and the β-catenin nuclear translocation^157^. Given the ability of the Wnt/β-catenin/TCF4 pathway in repressing HIV transcription, its targeting in “*shock and kill strategies*” deserve further investigations.

This study has several limitations. The effect of ATRA was studied on bulk MDMs. Given the functional particularities of CD16^+^ *versus* CD16^+^ monocytes in terms of RA receptor alpha (RARA) expression, RALDH activity^70,71^ and HIV-1 replication/dissemination^18,19^, the capacity of ATRA to selectively modulate HIV-1 replication in these subsets remains to be investigated. While the functional characterization of CD4+ T-cells carrying VR generated valuable insights into the host-cell determinants associated with integrative HIV-1 infection^158^, such studies are needed for macrophages as well, especially in the context of a tremendous heterogeneity at single-cell level^159^. This manuscript provides original evidence that ATRA blocks the antiviral activity of SAMHD1 *via* phosphorylation, but the molecular mechanism by which antiviral response are blunted by ATRA remain to be elucidated. Of particular notice, pioneering work by Blalock *et al.* demonstrated that RA transcriptionally represses interferon production^160^. Finally, although our RNA Sequencing results reveled a down regulation of estrogen receptor 1 (ESR1) by ATRA in MDMs (<u>Supplemental File 2</u>), in this study we did not address sex-related differences in the capacity of RA to modulate HIV-1 replication in macrophages. Given the reported interplay between RA and estrogen pathways^161^, such studies become highly relevant.

In conclusion, our results demonstrate that HIV-1 efficiently replicates in ATRA-exposed MDMs *via* mTOR-dependent mechanisms that involve ***i)*** an increase in CCR5 expression with the selection of R5 HIV-1 strains; ***ii)*** a reduced SAMHD1-mediated restriction at the level of reverse transcription; ***iii)*** a sustained transcription in part explained by a decrease in the expression/activation of the Wnt/β-catenin pathway; and ***iv)*** a more efficient translation potentially *via* a reduced INSIG1-mediated Gag degradation (<u>Graphical Abstract</u>). These findings point to mechanisms that may be targeted in macrophages, as well as in CD4^+^ T-cells, to limit or boost HIV-1 replication. Although short-lived MDMs may not contribute to VR persistence in ART-treated PLWH, in the intestinal environment rich in RA, these MDMs likely contribute to the establishment of primary HIV-1 infection and viral rebound upon ART interruption. Therefore, HIV eradication strategies should consider MDMs as therapeutic targets.

## Authors’ contributions

JD designed and performed research/experiments, analyzed data, prepared figures, and wrote the manuscript. AC designed and performed research, analyzed data, prepared figures, and contributed to manuscript revisions. JPG analyzed the RNA Sequencing data, prepared figures, and contributed to manuscript revisions. LRM, AF, TRWS, CDNY, EMG, and ERC performed experiments and/or provided reagents/protocols, and contributed to manuscript revisions. JPR provided access to clinical samples/information, set up clinical research protocols, and contributed to manuscript revision. PA conceived the research study hypothesis, designed the research, analyzed results, prepared the figures, and wrote the manuscript. All co-authors approved the submission of this manuscript.

## Supporting information

Supplemental Figure gels Western blot

## Acknowledgments

The authors thank Dr. Dominique Gauchat, Philippe St Onge, and Dr. Gael Duluth (Flow Cytometry Core Facility, CHUM-Research Center, Montréal, QC, Canada) for expert technical support with polychromatic flow cytometry sorting; Olfa Debbeche and Laurent Knaffo (Biosafety Level 3 Core Facility CHUM-Research Cente, Montréal, QC, Canada); Mario Legault (FRQ-S/AIDS and Infectious Diseases Network; Montréal, QC, Canada) for help with ethical approvals and informed consents; Josée Girouard and Angie Massicotte (McGill University Health Centre, Montréal, QC, Canada) for their key contribution to blood collection and clinical information from PLWH and uninfected study participants. The authors also thank Dr. Dana Gabuzda (Dana-Farber Cancer Institute, Boston, Massachussets, USA), Dr Roger J Pomerantz (Thomas Jefferson University, Philadelphia, Pennsylvania, USA) and Dr Michel Tremblay (Université Laval, Quebec, QC, Canada) for providing us with VSV-G and HIV plasmids. Finally, the authors acknowledge the key contribution of all study participants for their precious gift of leukapheresis essential for this study.

## STAR METHODS

## RESOURCE AVAILABILITY

### Lead contact

Further information and requests for resources and reagents should be directed to and will be fulfilled by the lead contact Petronela Ancuta.

### Materials availability

This study did not generate unique reagents.

### Data and code availability

RNA-seq data have been deposited at Gene Expression Omnibus (GEO) database under accession GSE226653 and are publicly available as of the date of publication, as listed in the Key Resource Table. Original western blot images have been deposited at Mendeley database (https://data.mendeley.com).

Any additional information required to reanalyze the data reported in this paper is available from the lead contact upon request.

This paper does not report original code.

## EXPERIMENTAL MODEL AND STUDY PARTICIPANT DETAILS

### Study participants

Leukapheresis samples were collected from HIV-uninfected individuals (HIV^-^), as we previously described^71^. PBMC from leukapheresis were isolated by gradient centrifugation using the lymphocyte separation medium (Wisent, Saint-Jean-Baptiste/Canada), and preserved frozen in 10% DMSO (SIGMA, St. Louis/United States) in fetal bovine serum (FBS; Wisent, Saint-Jean-Baptiste/Canada) until use.

### Ethics statement

A written informed consent following the guidelines of the Declaration of Helsinki and approved by the Institutional Ethics Review Board of the McGill University Health Center (MUHC; Montréal, Québec, Canada) and the Centre de Recherche du Centre Hospitalier de l’Université de Montréal (CR-CHUM; Montréal, Québec, Canada) was provided, clarified, and signed by all participants in this study.

### Monocyte isolation

PBMCs from HIV^-^ individuals were used to isolate monocytes by negative selection using a pan monocyte magnetic associated cell sorting (MACS) isolation kit (Miltenyi, Bergisch Gladbach/Germany) using a MACS buffer FACS buffer (1X PBS, 10% FBS, 2 mM EDTA). Typically, monocyte purity was >95 %, with less than 1% contaminations in CD3^+^CD4^+^ T-cells, as determined by flow cytometry upon staining with appropriate Abs (Supplemental Figure 1), as we previously reported^71^.

### Generation of monocyte-derived macrophages

MDMs were generated by culturing monocytes in 48-well plates (Costar, Arizona/United States) (10^6^ monocytes/well/ml) in the presence of M-CSF for 6 days. The MDMs differentiation media consisted of RPMI-1640 (Thermo Fisher; Waltham/United States), 10% of FBS, 1% penicillin/streptomycin (Thermo Fisher, Waltham/United States), and M-CSF (20 ng/mL; R&D Systems, Minneapolis/United States). Media containing M-CSF was refreshed every 2 days. In parallel, MDMs were generated in the presence of ATRA (Sigma, St. Louis/United States) and exposed or not to the following drugs: INK128 (an MTORC1/2 inhibitor; Cayman Chemical, Ann Arbor/United States) and/or PRI-724 (a Wnt/β-catenin inhibitor; Selleck, Houston/United States) and/or PNU-74654 (a Wnt/β-catenin inhibitor; Selleck, Houston/United States). All drugs were titrated for their effect on MDMs viability and optimal concentrations were used, as indicated in the Figure legends.

### Flow cytometry analysis

Surface staining was performed on PBMC and monocytes, as well as on MDMs harvested from 48-well plates using cold 1X PBS. Cells were washed using a FACS buffer (1X PBS, 10% FBS, EDTA 2mM, 0.2% sodium azide) and incubated with the following antibodies: CD3, CD4, CD16, CD14, CD1c, HLA-DR, CD14, CD16, CD195/CCR5, CD184/CXCR4 (Supplemental Table 1). Intracellular staining of MDMs infected with HIV-1 *in vitro* was performed with the HIV-p24 Abs (KC57; Beckman Coulter, Brea/United States) was performed using the BD cytofix/cytoperm fixation/permeabilization solution kit (BD Biosciences, Franklin Lakes/United States) according to the manufacturer’s protocols. To exclude dead cells, the live/dead fixable aqua dead cell stain kit (Thermo Fisher, Waltham/United States) was used. In addition, positive gates were placed according to fluorescence minus one (FMO) strategy, as previously reported^162^. Samples were acquired by flow cytometry using the LSRIIA cytometer and the BD FACS Diva software (BD Bioscience, San Jose, CAL; California). Finally, results were analyzed using BD Flowjo (Tree Star, Inc., Ashland, Oregon, USA).

### RNA extraction and mRNA expression by real-time RT-PCR

Dual DNA/RNA extraction was performed using the AllPrep DNA/RNA/miRNA universal kit (Qiagen; Hilden/Germany), according to the manufacturer’s protocol. Briefly, RLT plus buffer containing β-mercaptoethanol was added directly onto the MDMs monolayer to lyse the cells. Each lysed sample was transferred onto an AllPrep mini spin column. RNA was eluted at a final volume of 30 µL. The expression of HIC1, TCF4, CCND2, CTNNB1, GSK3β and PPARγ mRNA was quantified using a one-step SYBR green real-time PCR kit (Qiagen, Hilden/Germany) and the Lightcycler 480 II (Roche, Basel/Switzerland), using QuantiTect primers (Qiagen, Hilden/Germany). The control 28S rRNA primers (were designed as we previously reported ^71^ (Supplemental Table 1) and purchased from Integrated DNA Technologies (IDT, Newark/United States). Real-time RT-PCR was performed in triplicates on various quantities of total RNA for the quantification of PPARγ (70 ng), HIC1 (50 ng), TCF4 (25 ng), CCND2 (10 ng) CTNNB1 (10 ng), GSK3β (10 ng) mRNA, and 28S rRNA (2 ng). Negative controls (without RNA, without RT enzyme, without mix) were performed for each transcript and the mRNA expression was normalized relative to the 28S rRNA. mRNA expression levels were calculated based on the relative differences (delta CT) between the target gene of interest (PPARγ, HIC1, TCF4, CCND2, CTNNB1, GSK3β) and the control gene (28S rRNA), as we previously reported^70^.

### HIV-1 infection *in vitro*

The X-tremeGENE^TM^ HP DNA transfection reagent (Roche, Basel/Switzerland) was used to generate HIV-1 stocks by transfecting 293T cells with plasmids obtained from the National Institute of Health (NIH) AIDS Research Program. The following molecular clones were used in this study: *i)* replication-competent R5 NL4.3BaL (HIV_NL4.3BaL_), *ii)* T/F THRO (HIV_THRO_), *iii)* replication-competent X4 NDK and *iv)* replication-defective VSV-G-pseudotyped HIV (HIV_VSV-G_) (env-deficient NL4.3 provirus pseudotyped with the VSV-G envelope and expressing *gfp* in place of *nef*). Viral stocks were quantified by HIV-p24 ELISA and titrated on TCR-activated CD4^+^ T-cells for the identification of optimal infectious concentrations. For infection, MDMs plated in 48-well plates at a density of 10^6^ cells/well were exposed to HIV-1 (30 ng HIV-p24/300 µL/well) and incubated at 37 °C for three hours. Unbound virions were removed by three-times washing with 1 mL media/well. MDMs were further cultured for up to fifteen days in media containing M-CSF (10 ng/ml) in the presence/absence of ATRA and/or other drugs, as indicated in Figure legends. The media containing M-CSF and/or drugs was refreshed every three days post-infection. Cell-culture supernatants were used for HIV-p24 quantification by ELISA. In parallel, MDMs were collected for real-time PCR quantification HIV-DNA levels and flow cytometry analysis of intracellular HIV-p24 expression at day 3 and 15 post-infection, respectively

### Quantification of early HIV-1 reverse transcripts by SYBR Green PCR

MDMs were digested directly in 48-well plates in a lysis buffer containing 0.1mM Tris HCl pH 8.0, 0.5% Tween® 20 detergent, 10 mg/mL proteinase K (Thermo Fisher, Waltham/United States), and molecular grade water (Wisent, Saint-Jean-Baptiste/Canada) at a concentration of 50,000 cells/15 μL (or 200 μL per 48-well plate). The quantification of different forms of HIV-DNA was performed using nested real-time PCR, relative to CD3 as a housekeeping gene, using primers/SYBR Green as indicated in Supplemental Table 2, as we previously described ^73,76^. Serial dilutions from 3x10^5^ to three ACH-2 T-cells were used as a standard curve for early reverse transcripts quantification. Amplification products from the first PCR reaction were diluted by a factor of ten before adding buffer, primers, and SYBR Green. The limit of detection for this assay is three HIV/CD3 copies per test. All PCR reactions were performed in triplicates

### Quantification of late HIV-1 reverse transcripts and integrated HIV-DNA by nested real-time PCR

MDMs lysates were prepared as described above. The quantification of Gag and integrated HIV-DNA was performed using nested real-time PCR, relative to CD3 as a housekeeping gene, using primers/probes as indicated in Supplemental Table 2, as we previously described^73,76^. Briefly, late reverse transcripts were quantified using primers directed against Gag HIV-1 regions (45 Amplification cycles), and integrated HIV-DNA levels (45 Amplification cycles) were quantified using primers against the Alu repetitive sequences and HIV-LTR region^158,163^. Serial dilutions from 3x10^5^ to three ACH2 cells were used as a standard curve for late reverse transcripts and HIV-DNA quantification. Amplification products from the first PCR reaction were diluted by a factor of ten before adding buffer, primers, and probes. The limit of detection for this assay is three HIV/CD3 copies per test. All PCR reactions were performed in triplicates

### Quantification of HIV-p24 by ELISA

The HIV-p24 levels were quantified in cell culture supernatants using a homemade ELISA assay, as we previously reported^71,76^.

### Illumina RNA Sequencing and analysis

The RNA sequencing and analysis was performed, as we recently reported^83^. Briefly, total RNA was extracted from ATRA-MDMs and DMSO-MDMs harvested prior HIV infection (Figure 1A) using AllPrep DNA/RNA/miRNA universal kit (Qiagen; Hilden/Germany), according to the manufacturer’s protocol. Genome-wide RNA sequencing profiles were generated by Genome Québec (Montreal, Québec, Canada) using the Illumina RNA-Sequencing technology (NovaSeq6000 S4 PE 100bp 25M reads). The paired-end sequencing reads were aligned to coding and non-coding transcripts from Homo Sapiens database GRCh 37 version75 and quantified with the Kallisto software version 0.44.0. The entire RNA-Sequencing data set and the technical information requested by Minimum Information About a Microarray Experiment (MIAME) are available at the GEO database under accession GSE226653. Statistical analyses were performed using R version 4.21. Differential expression analysis was performed using the limma Bioconductor R package (version 3.52.2) on the log2-counts per million (logCPM) transformed transcript-level and gene-level data. Differentially expressed genes (DEG) were identified based on p-values (p<0.05), adjusted p-values (adj. p<0.05) and fold-change (FC, cutoff 1.3) (<u>Supplemental Files 1-2</u>). Gene set variation analysis (GSVA; C2, C3, C5, C7, C8, and Hallmark databases; https://www.gsea-msigdb.org/gsea/msigdb/index.jsp) was performed using the GSVA method (package version 1.344.2) on the logCPM data using a Gaussian cumulative distribution function. Finally, genes presenting RARE in their promoters were identified using the ENCODExplorerData (version 0.99.5) and listed in <u>Supplemental File</u> <u>3</u>.

### Western blotting

Total lysates of MDMs (3x10^6^ per condition) were generated using the Radio-immunoprecipitation Assay (RIPA) Buffer 1X (Cell Signaling, Danvers/United States) containing phosphatase (PhosSTOP; Roche, Basel/Switzerland) and protease inhibitors (Complete, Mini, EDTA-free protease inhibitor; Roche, Basel/Switzerland). Total protein content from each condition was quantified using DC Protein Assay (Bio-Rad, Hercules/United States) in triplicate samples. SDS-PAGE gel electrophoresis was performed on a gradient polyacrylamide gels for 75 minutes at 130 volts and transferred on immobilon-PSQ polyvinylidene difluoride (PVDF) membranes (Sigma, St. Louis/United States) for 75 minutes at 100 volts. PVDF membranes were blocked with Tris-Buffered Saline (TBS) 0.1% Tween 5% Bovine Serum Albumin (BSA) for 45 minutes and incubated overnight with primary antibodies against target proteins at 4 °C (Supplemental Table 1). PVDF membranes were washed four times with TBS 0.1% Tween and incubated with HRP-linked secondary Abs (Supplemental Table 1) for one hour at room temperature. All Abs were diluted with blocking TBS 0.1% Tween 5% BSA. PVDF membranes were washed four times and proteins were revealed with chemiluminescence western blotting substrates (Bio-Rad, Hercules/United States). PVDF membranes were reused by using a reblot stripping solution (Sigma St. Louis/United States). A Chemidoc imaging system from Bio-Rad was used for chemiluminescence and colorimetric detection to visualize the bands on PVDF membranes and Image lab software (Sigma St. Louis/United States) was used to quantify the band intensity between each condition and each donor.

## QUANTIFICATION AND STATISTICAL ANALYSIS

### Statistical analysis

Statistical analyses were performed using the GraphPad Prism 9 software (GraphPad Software, Inc.). To determine the statistical significance between two matched groups, the Wilcoxon test was used. P-values <0.05 were considered statistically significant. The p-values are indicated in all Figures.

