## Supplemental Figures 1-8 for "Retinoic Acid Boosts HIV-1 Replication in Macrophages *via* CCR5/SAMHD1-Dependent and mTOR-Modulated Mechanisms"

### Slide 1
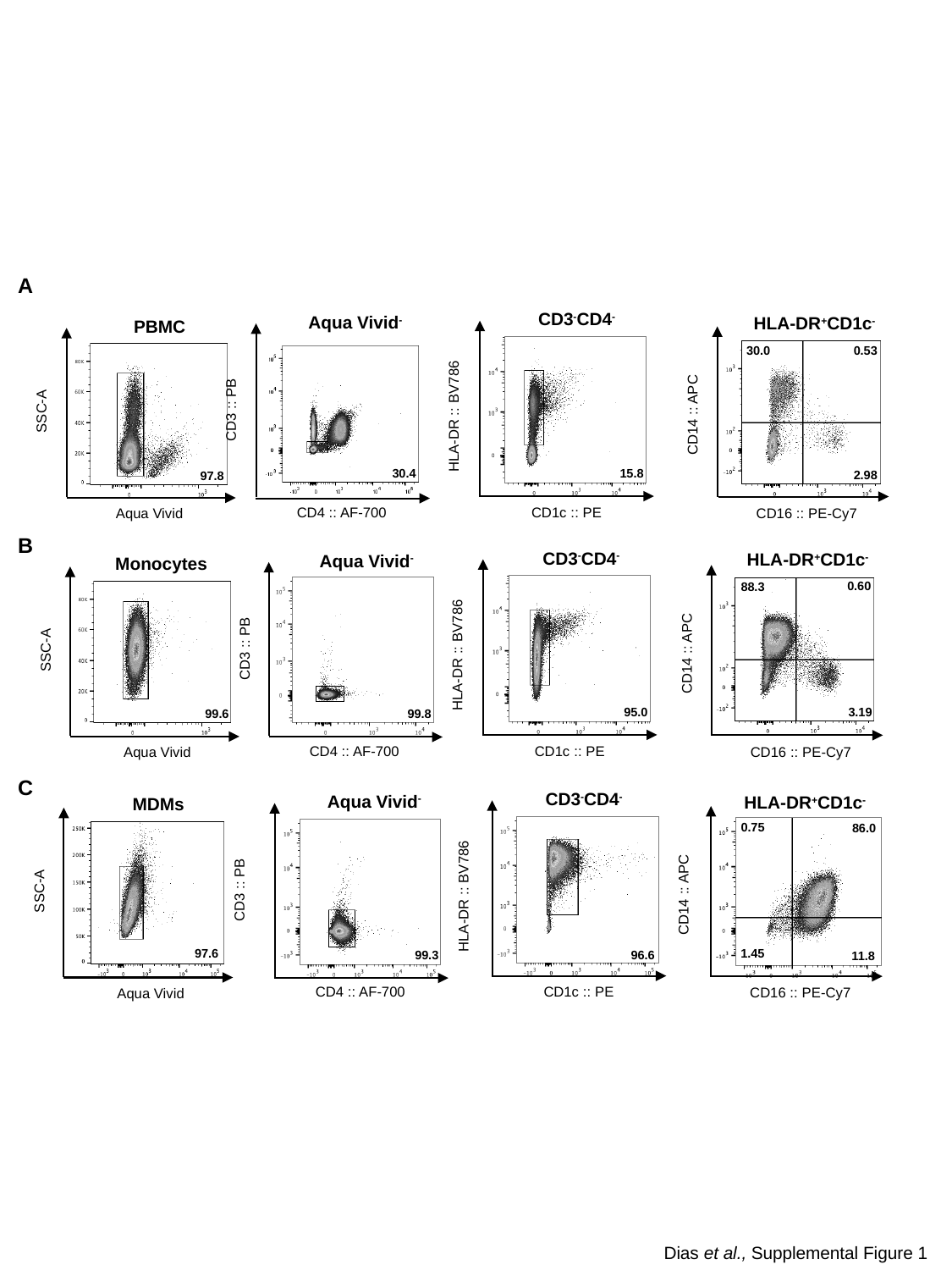

A
CD3^-^CD4^-^
HLA-DR :: BV786
CD1c :: PE
15.8
Aqua Vivid^-^
CD3 :: PB
CD4 :: AF-700
HLA-DR^+^CD1c^-^
CD14 :: APC
CD16 :: PE-Cy7
30.0
0.53
2.98
PBMC
SSC-A
Aqua Vivid
97.8
30.4
B
CD3^-^CD4^-^
HLA-DR :: BV786
CD1c :: PE
95.0
HLA-DR^+^CD1c^-^
CD14 :: APC
CD16 :: PE-Cy7
0.60
88.3
3.19
Aqua Vivid^-^
CD3 :: PB
CD4 :: AF-700
99.8
Monocytes
SSC-A
Aqua Vivid
99.6
C
CD3^-^CD4^-^
HLA-DR :: BV786
CD1c :: PE
96.6
Aqua Vivid^-^
CD3 :: PB
CD4 :: AF-700
99.3
HLA-DR^+^CD1c^-^
CD14 :: APC
CD16 :: PE-Cy7
86.0
0.75
1.45
11.8
SSC-A
Aqua Vivid
97.6
MDMs
Dias et al., Supplemental Figure 1

### Slide 2
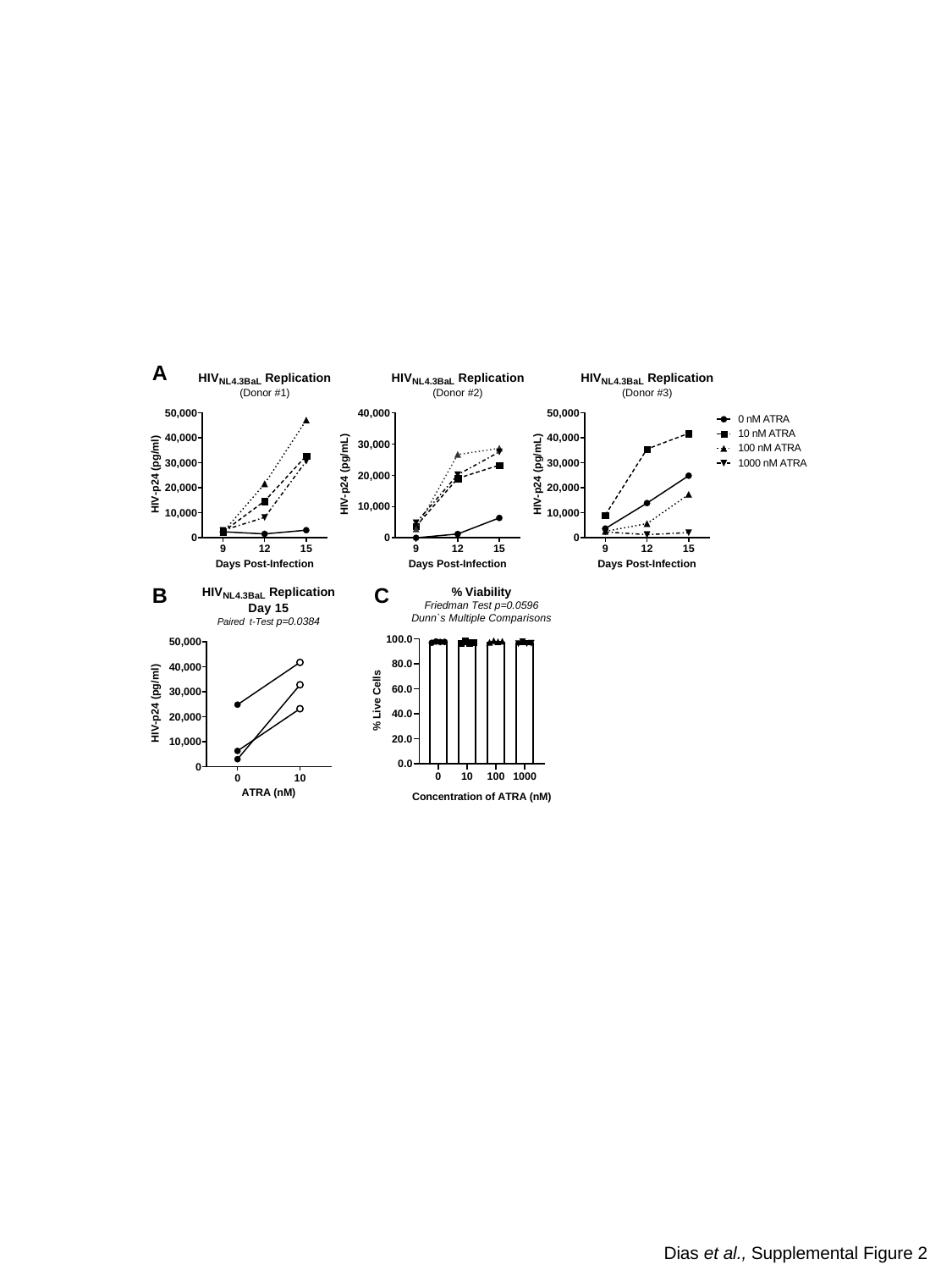

A
B
C
Dias et al., Supplemental Figure 2

### Slide 3
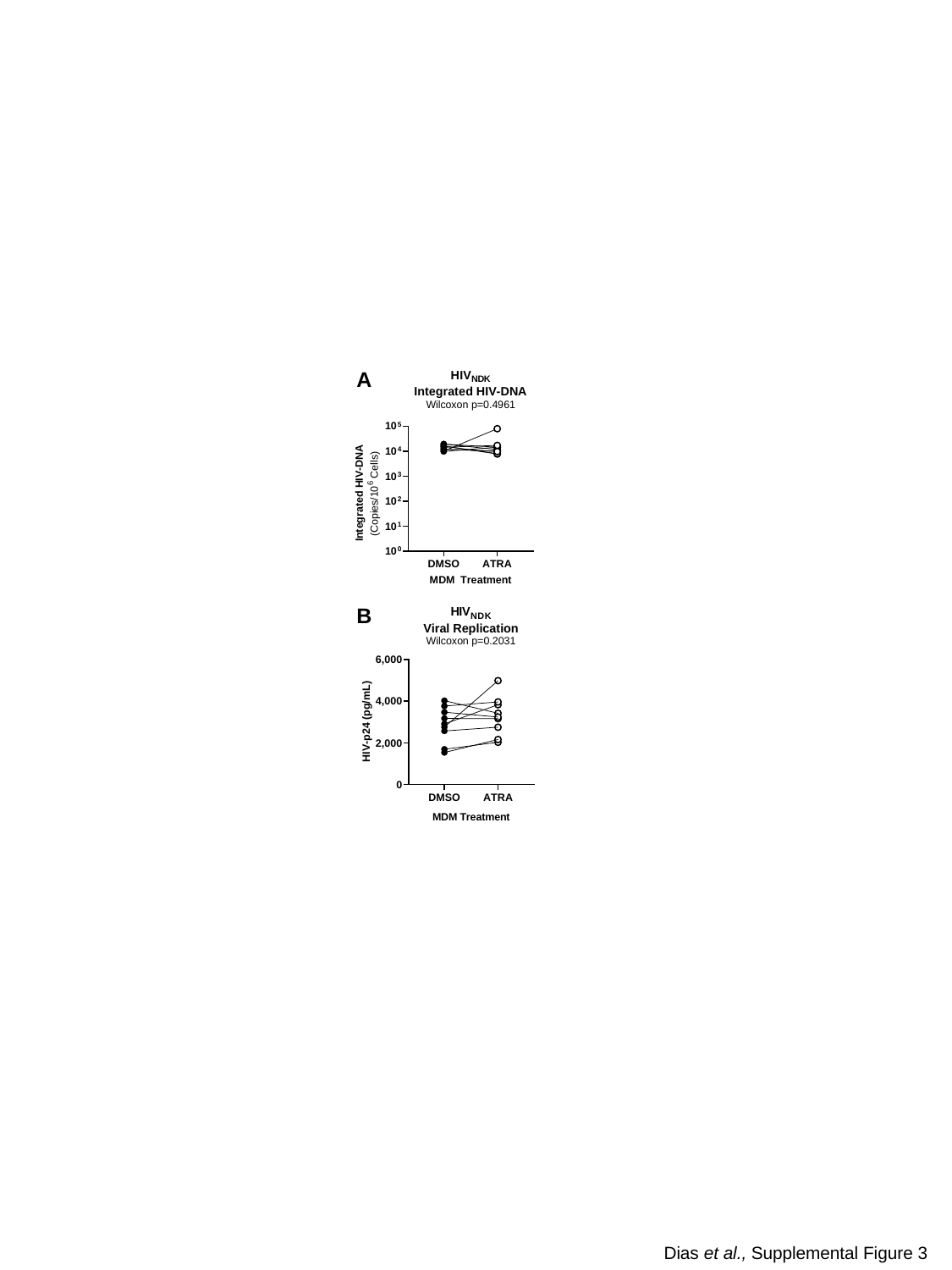

A
B
Dias et al., Supplemental Figure 3

### Slide 4
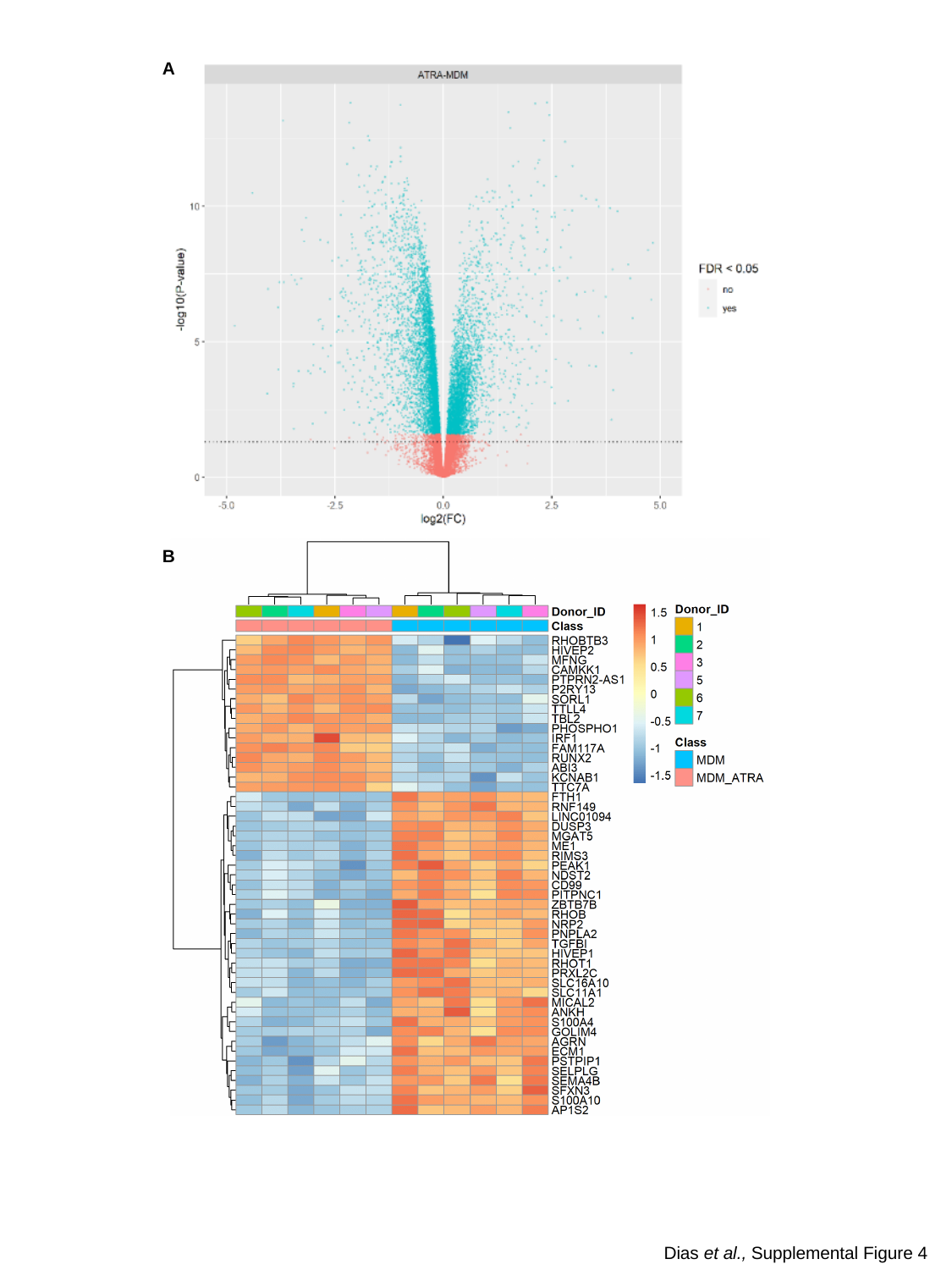

A
B
Dias et al., Supplemental Figure 4

### Slide 5
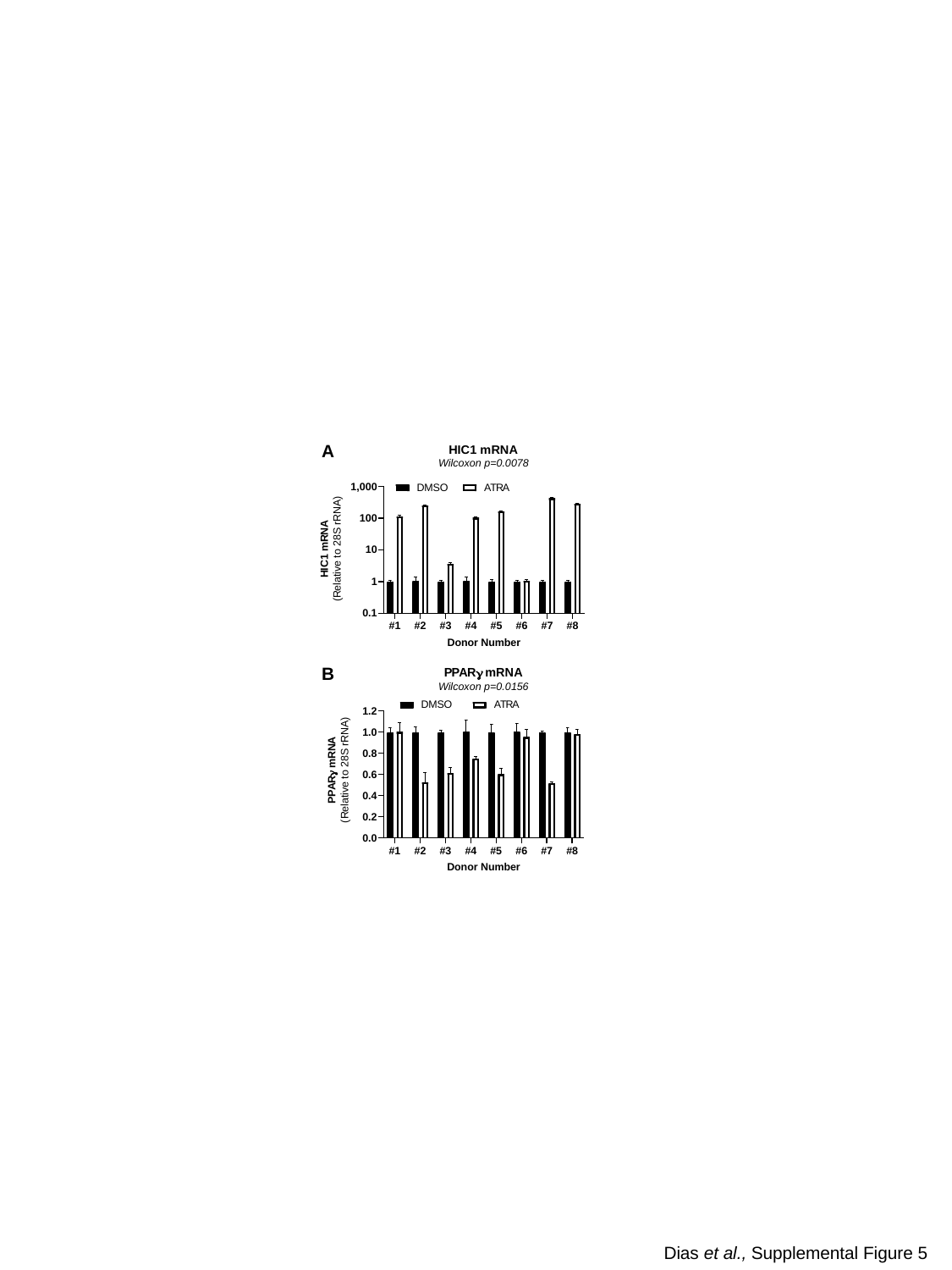

A
B
Dias et al., Supplemental Figure 5

### Slide 6
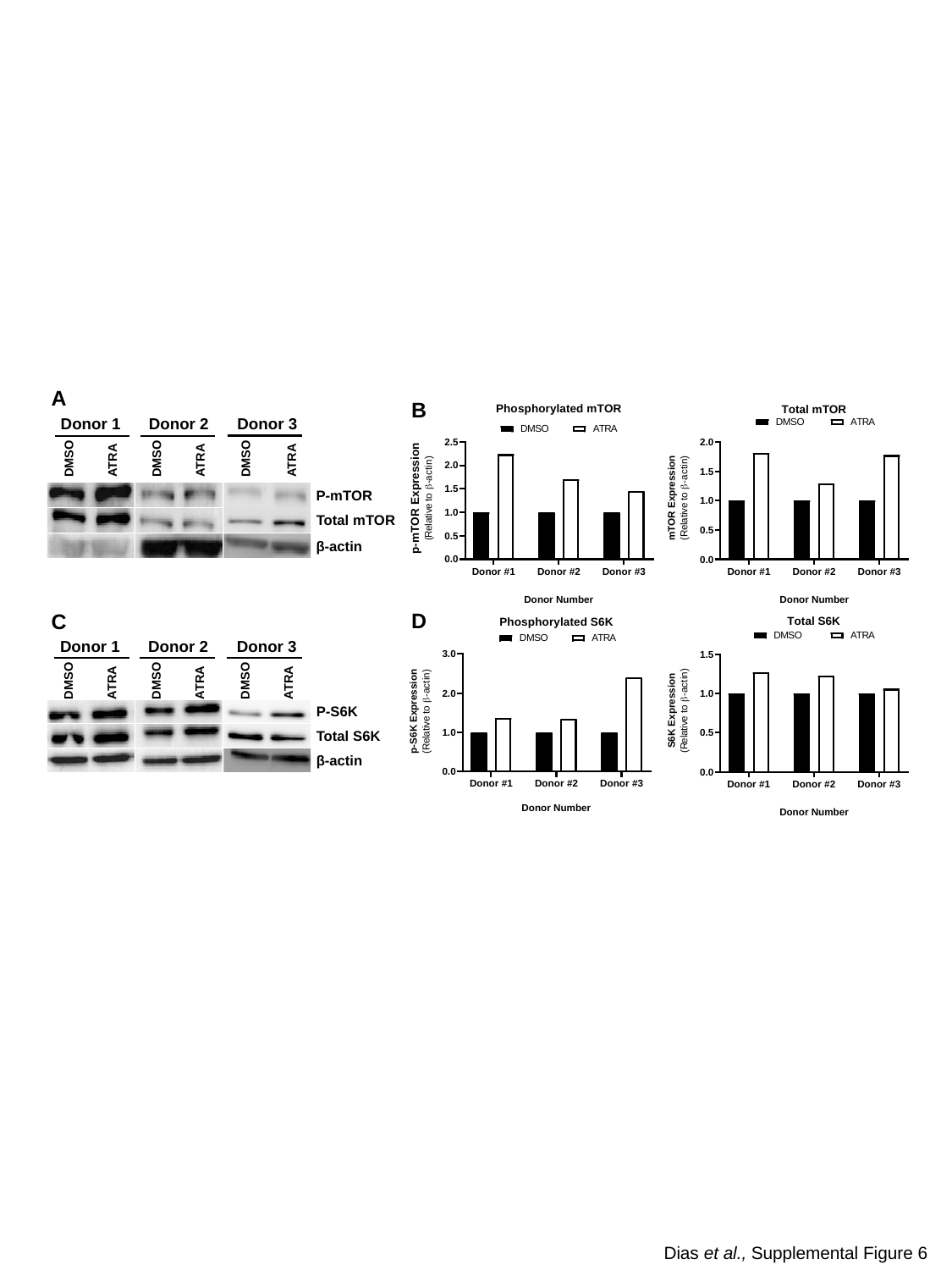

A
B
Donor 3
Donor 1
Donor 2
DMSO
DMSO
DMSO
ATRA
ATRA
ATRA
P-mTOR
Total mTOR
β-actin
D
C
Donor 3
Donor 1
Donor 2
DMSO
DMSO
DMSO
ATRA
ATRA
ATRA
P-S6K
Total S6K
β-actin
Dias et al., Supplemental Figure 6

### Slide 7
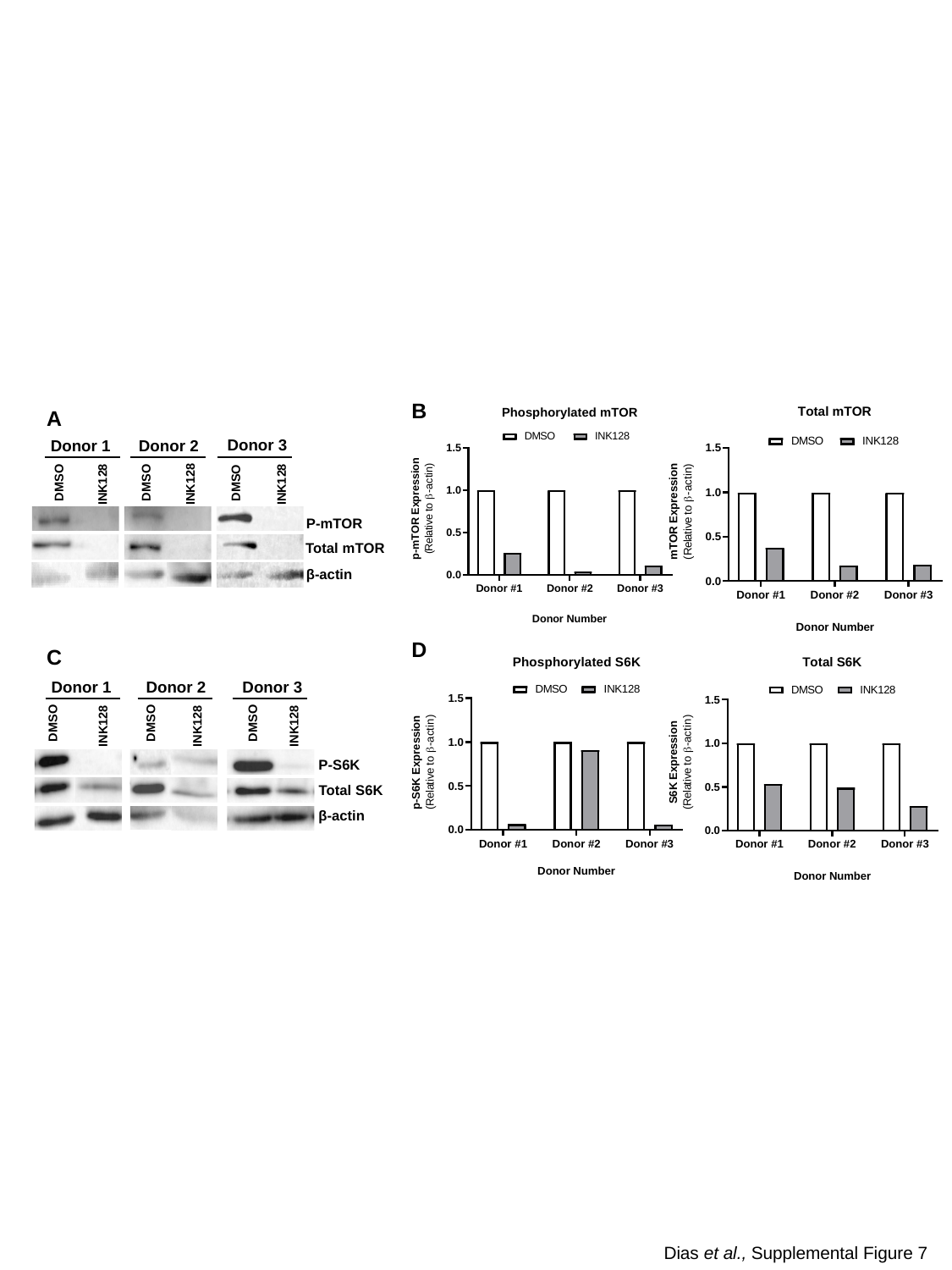

B
A
Donor 3
Donor 1
Donor 2
DMSO
DMSO
DMSO
INK128
INK128
INK128
P-mTOR
Total mTOR
β-actin
D
C
Donor 3
Donor 1
Donor 2
DMSO
DMSO
DMSO
INK128
INK128
INK128
P-S6K
β-actin
Total S6K
Dias et al., Supplemental Figure 7

### Slide 8
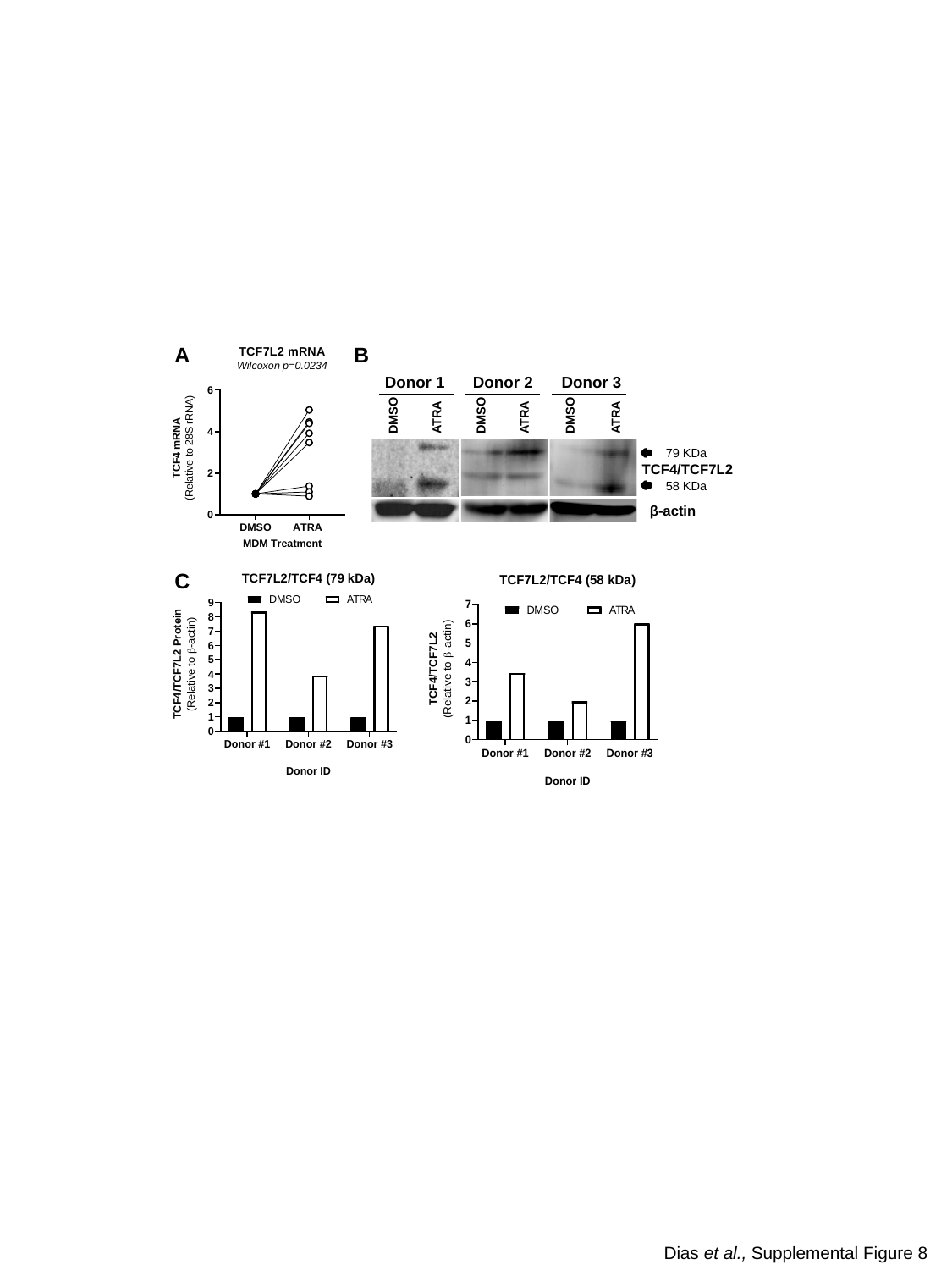

A
B
Donor 3
Donor 1
Donor 2
DMSO
DMSO
DMSO
ATRA
ATRA
ATRA
79 KDa
TCF4/TCF7L2
58 KDa
β-actin
C
Dias et al., Supplemental Figure 8
