## Supplemental Table 1 for "Retinoic Acid Boosts HIV-1 Replication in Macrophages *via* CCR5/SAMHD1-Dependent and mTOR-Modulated Mechanisms"

**Key Resources Table 1**

|  | SOURCE | IDENTIFIER |
| --- | --- | --- |
| **Leukapheresis** | | |
| Lymphocyte Separation Medium (LSM) | Wisent | Cat#305-010-CL |
| Fetal Bovine Serum (FBS) | Wisent | Cat#091-150 |
| Trypan Blue | Thermo Fisher | Cat#15250061 |
| Dimethyl Sulfoxide (DMSO) | Sigma | Cat#34869-500mL |
| RPMI 1640 Medium (RPMI) | Thermo Fisher | Cat#11875119 |
| **Monocyte isolation** | | |
| Pan Monocyte Isolation Kit | Miltenyi | Cat#130-096-537 |
| Trypan Blue | Thermo Fisher | Cat#15250061 |
| RPMI 1640 Medium (RPMI) | Thermo Fisher | Cat#11875119 |
| Penicillin/streptomycin (PenStep) | Thermo Fisher | Cat#15140122 |
| Fetal Bovine Serum (FBS) | Wisent | Cat#091-150 |
| Phosphate Buffered Saline (PBS) | Thermo Fisher | Cat#10010023 |
| MACS LS Columns | Miltenyi | Cat# 130-042-401 |
| Pre-Separation Filters 30 µm | Miltenyi | Cat# 130-041-407 |
| EDTA | Bioshop | Cat#EDT001.1 |
| **Culture conditions** | | |
| RPMI 1640 Medium (RPMI) | Thermo Fisher | Cat#11875119 |
| All-trans Retinoic Acid (ATRA) | Sigma | Cat#R2625-50MG |
| Sapanisertib (INK128) | Cayman Chemical | Cat#11811-1 |
| PRI-724 (ICG-001) | Selleck | Cat# S8968 |
| PNU-74654 | Selleck | Cat# S8429 |
| Recombinant Human Macrophage Colony Stimulating Factor (M-CSF) | Cedarlane | 216-MC-025 |
| Dimethyl Sulfoxide (DMSO) | Sigma | Cat#34869-500mL |
| **Flow cytometry** | | |
| Mouse anti-human CD3 Pacific Blue (Clone UCHT1) | BD | Cat#558117 |
| Mouse anti-human CD4 Alexa Fluor 700 (Clone RPA-T4) | BD | Cat#557922 |
| Mouse anti-human CD16 Phycoerythrin-Cyanine 7 (Clone 3G8) | BD | Cat#560918 |
| CD14 | BD | Cat#555399 |
| Anti-human CD1c (BDCA-1) Phycoerythrin (Clone REA694) | Miltenyi | Cat#130-110-536 |
| Anti-human HLA-DR Brilliant Violet 785 (Clone L243) | Biolegend | Cat#307642 |
| Mouse Anti-human CD195 Fluorescein Isothiocyanate (Clone 2D7/CCR5) | BD | Cat#555992 |
| CD184 CXCR4 Allophycocyanine (Clone 12G5) | Biolegend | Cat#306510 |
| HIV-1 core (p24) antigen-RD1 (Clone KC57) | Beckman Coulter | Cat#6604667 |
| Fixation/Permeabilization Kit | BD | Cat#554714 |
| LIVE/DEAD Fixable Aqua Dead Cell Stain Kit (405 nm excitation) | Thermo Fisher | Cat#L34957 |
| Phosphate Buffered Saline (PBS) | Thermo Fisher | Cat#10010023 |
| Fetal Bovine Serum (FBS) | Wisent | Cat#091-150 |
| Sodium Azide | Bioshop | Cat#SAZ001.250 |
| GSK3B pS9 Antibody, anti-human/mouse/rat REAfinity APC | Miltenyi | Cat#130-106-901 |
| Beta Catenin Monoclonal Antibody (15B8), Alexa Fluor 488 | Invitrogen | Cat#53-2567-41 |
| Formaldehyde solution 37 wt. % in H2O | Sigma | Cat#F1635-500ML |
| **Western blot** | | |
| Radioimmunoprecipitation Assay Buffer (RIPA) 10X | Cell Signaling | Cat#9806S |
| Phosphatase Inhibitor (PhosSTOP) | Roche | Cat#4906845001 |
| Complete, Mini, EDTA-free protease inhibitor | Roche | Cat#11836170001 |
| Detergent Compatible (DC) Protein Assay | Bio-Rad | Cat#5000111 |
| 30% Bis-acrylamide Solution | Bioshop | Cat#ACR010.500 |
| UltraPure Tris Buffer | Thermo Fisher | Cat#15504-020 |
| Sodium Dodecyl Sulfate (SDS) | Bio-Rad | Cat#1610302 |
| N,N,N’,N’-Tetramethyl Ethylenediamine (TEMED) | Sigma | Cat#8087420005 |
| Ammonium Persulfate (APS) | Bioshop | Cat#AMP001.100 |
| Immobilon-PSQ Polyvinylidene Difluoride (PVDF) | Sigma | Cat#ISEQ00010 |
| 4X Laemmli Sample Buffer | Bio-Rad | Cat#1610747 |
| 2-Mercaptoethanol | Sigma | Cat#M6250 |
| Precision Plus Protein Dual Color Standards | Bio-Rad | Cat#1610374 |
| Glycine | BioShop | Cat#GLN001.500 |
| Sodium Chloride | BioShop | Cat#SOD002 |
| Tween 20 | Fisher Scientific | Cat#BP337-500 |
| Bovine Serum Albumin (BSA) | BioShop | Cat#ALB001.500 |
| Phospho-mTOR (Ser2448) Antibody (1/1000) | Cell Signaling | Cat#2971S |
| mTOR (7C10) Rabbit mAb (1/1000) | Cell Signaling | Cat#2983 |
| Anti-phospho-ribosomal protein S6 Ser240/Ser244 (1/500) | EMD Millipore | Cat#07-2113 |
| S6 Ribosomal Protein (5G10) Rabbit mAb (1/1000) | Cell Signaling | Cat#2217 |
| TCF4/TCF7L2 (C48H11) Rabbit (1/1000) | Cell Signaling | Cat#2569 |
| Monoclonal Anti-β-actin Antibody Produce in Mouse (1/5000) | Sigma | Cat#A5441 |
| Anti-rabbit IgG HRP-linked Antibody (1/5000) | Cell Signaling | Cat#7074 |
| Goat Anti-Mouse IgG (H+L) Secondary Antibody, HRP (1/2000) | Thermo Fisher | Cat#32340 |
| Clarity and Clarity Max ECL Western Blotting Substrates | Bio-Rad | Cat#1705062S |
| ReBlot Plus Strong Antibody Stripping Solution | Sigma | Cat#2504 |
| Monoclonal anti-SAMHD1 mouse antibody clone OTI3F5 (30 µl) | Cedarlane | Cat#TA502024S |
| Polyclonal Anti-SAMHD1 (PHOSPHO THR592) (0,02 mg) antibody | Cedarlane | Cat#8005-0.02MG |
| HIC-1 Antibody (H-6) 200 µg/ml | santa cruz | Cat#sc-271499 |
| Methanol 99,98% | Fisher Scientific | Cat#BPA4084 |
| **Total DNA/RNA Extraction and mRNA Expression by RT-PCR** | | |
| All Prep DNA/RNA/miRNA Universal Kit | Qiagen | Cat#80224 |
| QuantiTect SYBR Green RT-PCR Kit | Qiagen | Cat#204243 |
| See Supplemental Table 1: Oligonucleotides | See Supplemental Table 1 : Oligonucleotides | See Supplemental Table 1 : Oligonucleotides |
| **Virus Strains** | | |
| HIV-1 THRO plasmid (pTHRO.c/2626), subtype B | NIH AIDS Reagent Program (Contribution of Dr. John Kappes and Dr. Christina Ochsenbauer) | Cat#11745 |
| NL4.3BaL HIV plasmid | From Dr. Dana Gabuzda (Dana-Farber Cancer Institute, Boston, MA, USA) | N/A |
| VSV-G Plasmid | National Institution of Health (NIH) | Cat#ARP-4693 |
| NL4.3BaLΔenv GFP | National Institution of Health (NIH | Cat#ARP-12637 |
| X-tremeGENE HP DNA Transfection Reagent | Roche | 6366244001 |
| Memory CD4+ T Cell Isolation Kit, human | Miltenyi | Cat#130-091-893 |
| 293T Cells | ATCC | Cat#CRL-3216 |
| DMEM High glucose, pruvate | Gibco | Cat#11995065 |
| Opti-MEM | Gibco | Cat#31985070 |
| Fetal Bovine Serum (FBS) | Wisent | Cat#091-150 |
| Phosphate Buffered Saline (PBS) | Thermo Fisher | Cat#10010023 |
| Trypsin EDTA with phenol red | Wisent | Cat#075-350 |
| **Early/Late Reverse Transcripts and Integrated HIV-DNA** | | |
| Tris HCl | BioShop | Cat#TRS002.500 |
| Tween 20 | Fisher Scientific | Cat#BP337-500 |
| Proteinase K | Fisher Bioreagents | Cat#25530-015 |
| Molecular Grade Water (H_2_O) | Wisent | Cat#809-115-CL |
| Thermus Aquaticus (TAQ) DNA Polymerase | Thermo Fisher | Cat#18038067 |
| Magnesium Chloride (MgCl_2_) Buffer | Thermo Fisher | Cat#18038067 |
| Deoxynucleoside Triphosphates (dNTP) | Thermo Fisher | Cat#10297018 |
| QuantiTect SYBR Green RT-PCR Kit | Qiagen | Cat#204243 |
| LC480 probe master mix | Roche | Cat#4707494001 |
| 10X PCR Buffer | Thermo Fisher | Cat#18038067 |
| Nonidet P-40 | Bioshop | Cat#NON505 |
| See Supplemental Table 2: Oligonucleotides | See Supplemental Table 2 : Oligonucleotides | See Supplemental Table 2 : Oligonucleotides |
| **ELISA** | | |
| Human IL-10 DuoSet Enzyme-linked Immunosorbent Assay (ELISA) Kit | RnD Systems | Cat#DY217B |
| p24 Enzyme-linked Immunosorbent Assay (ELISA) | Homemade. Hybridome provided by Dr. Michel J. Tremblay (Bounou et al., J Virol., 2002) | |
| Sodium Bicarbonate (NaHCO_3_) | Sigma | Cat#S5761 |
| Sodium Carbonate (Na_2_CO_3_) | Sigma | Cat#223530-500G |
| Thimerosal | Sigma | Cat#T5125-10G |
| Phosphate Buffered Saline (PBS) | Thermo Fisher | Cat#10010023 |
| Tween 20 | Fisher Scientific | Cat#BP337-500 |
| Triton X-100 | Sigma | Cat#X100-500mL |
| Trypan Blue | Thermo Fisher | Cat#15250061 |
| Bovine Serum Albumin (BSA) | BioShop | Cat#ALB001.500 |
| Streptavidin Horseradish Peroxidase (Strep-HRP) | Fisher Scientific | Cat#65R-S104PHRP |
| 3,3’,5,5’-Tetramethylbenzidine (TMB) | Quimigen | Cat#42R-TB10265R-S104PHRP |
| Phosphoric Acid (H_3_PO_4_) | Sigma | Cat#PX0996 |
| **Cell Lines** | | |
| HT-29 cell line | ATCC | Cat#HTB-38 |
| ACH-2 cell line | NIH HIV Reagent Program | Cat#ARP-349 |
| **Software** | | |
| FlowJo version 10 | BD | https://www.flowjo.com/ |
| GraphPad Prism 9 | GraphPad | https://www.graphpad.com/ |
| Image Lab | Bio-Rad | https://www.bio-rad.com/en-ca/product/image-lab-software?ID=KRE6P5E8Z |
