## Supplemental Table 2 for "Retinoic Acid Boosts HIV-1 Replication in Macrophages *via* CCR5/SAMHD1-Dependent and mTOR-Modulated Mechanisms"

| Identification | Sequence | Provider | Identifier |
| --- | --- | --- | --- |
| Primer AA55 | 5’-CGT CTA GAG ATT TTC CAC AC-3’ | IDT | N/A |
| Primer M667 | 5’-CTA ACT AGG GAA CCC ACT G-3’ | IDT | N/A |
| CD3 external primer 1 | 5’-ACT GAC ATG GAA CAG GGG AAG-3’ | IDT | N/A |
| CD3 external primer 2 | 5’- CCA GCT CTG AAG TAG GGA ACA TAT-3’ | IDT | N/A |
| Primer GagR | 5’- AGC TCC CTG CTT GCC CAT A-3’ | IDT | N/A |
| Primer Alu1 | 5’- TCC CAG CTA CTG GGG AGG CTG AGG-3’ | IDT | N/A |
| Primer Alu2 | 5’- GCC TCC CAA AGT GCT GGG ATT ACA G-3’ | IDT | N/A |
| Primer LM667 | 5’- ATG CCA CGT AAG CGA AAC TCT GGC TAA CTA GGG AAC CCA CTG-3’ | IDT | N/A |
| Primer LambdaT | 5’- ATG CCA CGT AAG CGA AAC T-3’ | IDT | N/A |
| Primer AA55M | 5’- GCT AGA GAT TTT CCA CAC TGA CTA A-3’ | IDT | N/A |
| Primer SK30 | 5’- GGT CTG AGG GAT CTC TAG-3’ | IDT | N/A |
| Primer SK29 | 5’- ACT AGG GAA CCC ACT GCT-3’ | IDT | N/A |
| CD3 internal primer 1 | 5’- CCT CTC TTC AGC CAT TTA AGT A-3’ | IDT | N/A |
| CD3 internal primer 2 | 5’- GGC TAT CAT TCT TCT TCA AGG T-3’ | IDT | N/A |
| Probe LTR-LC | 5’-LC640- CACTCAAGGCAAGCTTTATTGAGGC-3’-phosphate | TIB MolBiol | N/A |
| Probe LTR-FL | 5’- CACAACAGACGGGCACACACTACTTGA-3’-Flurescein | TIB MolBiol | N/A |
| Probe P1 | 5’-GGCTGAAGGTTAGGGATACCAATATTCCTGTCTC-3’-Flurescein | TIB MolBiol | N/A |
| Probe P2 | 5’-LC640- CTAGTGATGGGCTCTTCCCTTGAGCCCTTC-3’-phosphate | TIB MolBiol | N/A |
| Primer 28S Forward | 5′-CGAGATTCCTGTCCCCACTA-3′ | IDT | N/A |
| Primer 28S Reverse | 5′-GGGGCCACCTCCTTATTCTA-3′ | IDT | N/A |
| PPARγ | N.A. | Qiagen | GeneGlobal ID: QT00029841 |
| TCF7L2 | N.A. | Qiagen | GeneGlobal ID: QT00071120 |
| HIC1 | N.A. | Qiagen | GeneGlobal ID: QT00203175 |
| CTNNB1 | N.A. | Qiagen | GeneGlobal ID: QT00077882 |
| GSK3β | N.A. | Qiagen | GeneGlobal ID: QT00057134 |
| CCND2 | N.A. | Qiagen | GeneGlobal ID: QT00057575 |

**Supplemental Table 2: Primers and probes used for PCR and RT-PCR. Related to STAR Methods.**

*N.A, information not available; N/A, Not applicable*
