## Supplemental Figure legends 1-8 for "Retinoic Acid Boosts HIV-1 Replication in Macrophages *via* CCR5/SAMHD1-Dependent and mTOR-Modulated Mechanisms"

**Supplemental Figure 1: Phenotyping analysis of monocytes and MDMs.** Monocytes were isolated from PBMCs by negative selection using magnetic beads (Miltenyi). Monocyte-derived macrophages (MDMs) were generated by culturing monocytes in the presence of M-CSF (20 ng/mL) for six days, as depicted in Figure 1A. In each experiment, the purity of sorted monocytes and the phenotype of MDMs were determined by flow cytometry analysis of matched PBMCs **(A)**, monocytes **(B)**, and MDMs **(C)** upon staining with CD3, CD4, HLA-DR, CD1C, CD14 and CD16 Abs. The viability dye aqua Vivid was used to exclude dead cells from the analysis. Monocytes and macrophages were identified as cells with a CD3^-^CD4^-^HLA-DR^+^CD1c^-^ and CD3^-^CD4^-^HLA-DR^+^CD1c^-^CD14^+^CD16^+^ phenotype, respectively. Shown are results generated with cells from one donor representative of results generated with MDMs from 10 HIV-uninfected donors.

**Supplemental Figure 2: The dose-effect of ATRA on HIV-1 replication in MDMs.** MDMs were generated in the presence or the absence of different concentrations of ATRA (10, 100 and 1,000 nM), as depicted in Figure 1A, and were exposed to HIV_NL4.3BaL_ (30 ng HIV-p24/well per 10^6^ MDMs in 300 µl media) for 3 hours. Unbound virions were removed by extensive washing. Cells were cultured in media containing M-CSF (20 ng/mL), in the presence/absence of ATRA at the indicated concentrations. Cell-culture supernatants were collected every 3 days for HIV-p24 ELISA quantification, and fresh media containing M-CSF and/or ATRA was added every 3 days. **(A)** Shown are the kinetics of HIV-1 replication *per* donor and **(B)** statistical analysis performed at days 15 post-infection with MDMs from n=3 HIV-uninfected donors **(C)**. Cell viability was measured by flow cytometry using the Aqua Vivid viability dye (n=3). Paired t-Test **(B)** and Friedman and Dunn`s multiple comparison p-values **(C)** are indicated on the graphs.

**Supplemental Figure 3: ATRA does not impact CXCR4-tropic HIV-1 integration and replication in MDMs.** MDMs generated by culturing monocytes in media containing M-CSF (20 ng/mL), in presence or the absence of ATRA (10 nM), as in Figure 1A, were exposed to the CXCR4-troic (X4) HIV_NDK_ strain (50 ng HIV-p24/10^6^ cells in 300 µl). Shown are levels of (**A)** integrated HIV-DNA measured by real-time nested PCR and **(B)** HIV-p24 measured by ELISA in cell culture supernatants at day 3 post-infection. Experiments were performed with MDMs from n=8 HIV-uninfected individuals. Wilcoxon test p-values are indicated on the graphs.

**Supplemental Figure 4: Identification of genes modulated by ATRA in MDMs.** Total RNA extracted from MDMs generated in the presence (ATRA-MDMs) or the absence (DMSO-MDMs) of ATRA (10 nM) were used for genome-wide RNA sequencing (Illumina technology), as in Figure 3A. **(A)** The volcano plot depicts the log2 fold change (FC) in gene expression levels (x axis; cut-off: 1.3) and the log10 p-values for differentially expressed genes (DEG; y axis). **(B)** The heatmap depicts the top 50 upregulated and downregulated DEG in MDMs generated in ATRA-MDMs *versus* DMSO-MDMs. Experiments were performed with MDMs from n=6 HIV-uninfected individuals.

**Supplemental Figure 5: RT-PCR validation of differential PPARγ and HIC1 mRNA expression in ATRA-MDMs versus DMSO-MDMs.** Total RNA was extracted from MDMs generated in the presence or the absence of ATRA (10 nM), as in Figure 3. Shown are levels of PPARγ **(A)** and HIC1 mRNA expression **(B)**, as measured by SYBR-Green real-time RT-PCR. Wilcoxon p-values are indicated on the graphs. Experiments were performed with MDMs from n=8 HIV-uninfected individuals.

**Supplemental Figure 6: ATRA increases the expression and the phosphorylation of mTOR and S6K in MDMs.** MDMs were generated in the presence or the absence of ATRA (10 nM; Day -2 until Day 0), as in Figure 1A. Cells harvested prior HIV exposure were used to generated cell lysates that were subject to SDS-gel electrophoresis for the visualisation/quantification of total and phosphorylated/total mTOR expression **(A-B)**, as well as total/phosphorylated S6K expression **(C-D).** Shown is the visualisation of total mTOR (MW: 289 kDa), phosphorylated mTOR (MW: 289 kDa), total S6K (MW: 32 kDa) and phosphorylated S6K (MW: 32 kDa), and β-actin protein bands (MW: 42 kDa) **(A and C)** in MDMs upon incubation with specific Abs. Results in the graphs depict total (**right panels**) and phosphorylated (**left panels**) mTOR **(B)** and S6K expression levels **(D)** normalized to ß-actin levels. Experiments were performed with MDMs from n=3 HIV-uninfected individuals.

**Supplemental Figure 7: Exposure to INK128 decreases mTOR and S6K expression/phosphorylation in ATRA-treated MDMs.** Cell lysates from ATRA-MDMs exposed or not to INK128 for 48 hours were subject to SDS-gel electrophoresis for the visualisation of total/phosphorylated mTOR **(A-B)** and S6K expression **(C-D)** Shown is the visualisation of total mTOR, phosphorylated mTOR, total S6K and phosphorylated S6K, and β-actin protein bands **(A and C)** in ATRA-MDMs upon incubation with specific Abs. Results in the graphs depict total (**right panels**) and phosphorylated (**left panels**) mTOR **(B)** and S6K expression levels **(D)** normalized to ß-actin levels. Experiments were performed with MDMs from n=3 HIV-uninfected individuals.

**Supplemental Figure 8: ATRA modulates the TCF4 mRNA and protein expression.** ATRA-MDMs and DMSO-MDMs obtained as in Figure 3A, were used for the quantification of TCF4 mRNA expression by SYBR-Green RT-PCR **(A)**, as well as the visualisation/quantification of TCF4 protein expression by western blotting **(B-C).** Shown are statistical analysis of TCF4 mRNA expression in MDMs from n=8 HIV-uninfected donors **(A)**. Also, shown are TCF4 (MW: 79 KDa and 58 KDa) and β-actin (MW: 42 KDa) protein expression in MDMs **(B)** and relative expression of the two TCF4 bands (79 KDa, left panel; 58 KDa right panel) relative to β-actin **(C)** in MDMs from n=3 HIV-negative donors.
