## Supplemental Figure gels Western blot for "Retinoic Acid Boosts HIV-1 Replication in Macrophages *via* CCR5/SAMHD1-Dependent and mTOR-Modulated Mechanisms"

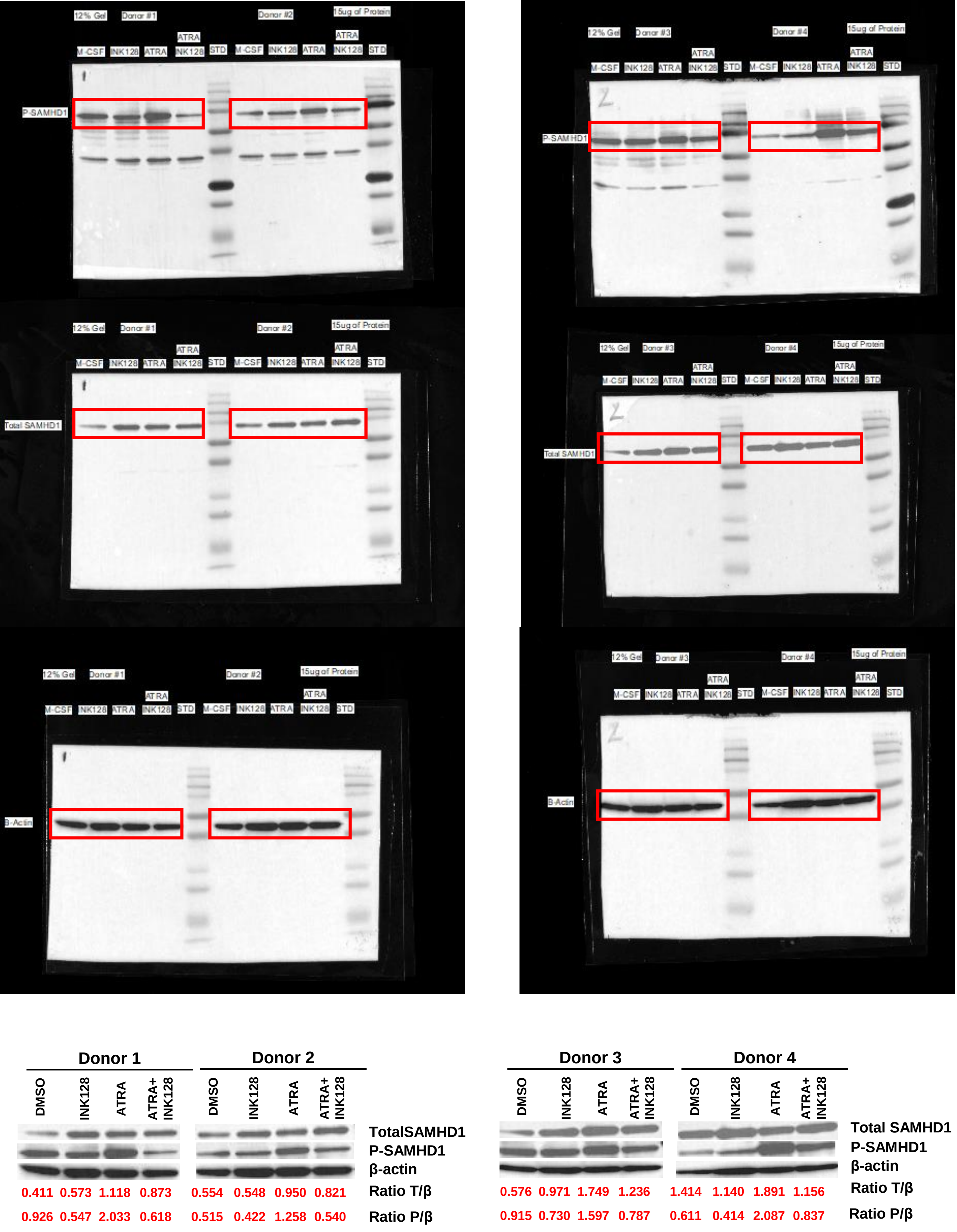


Dias *et al.*, Figure 6B

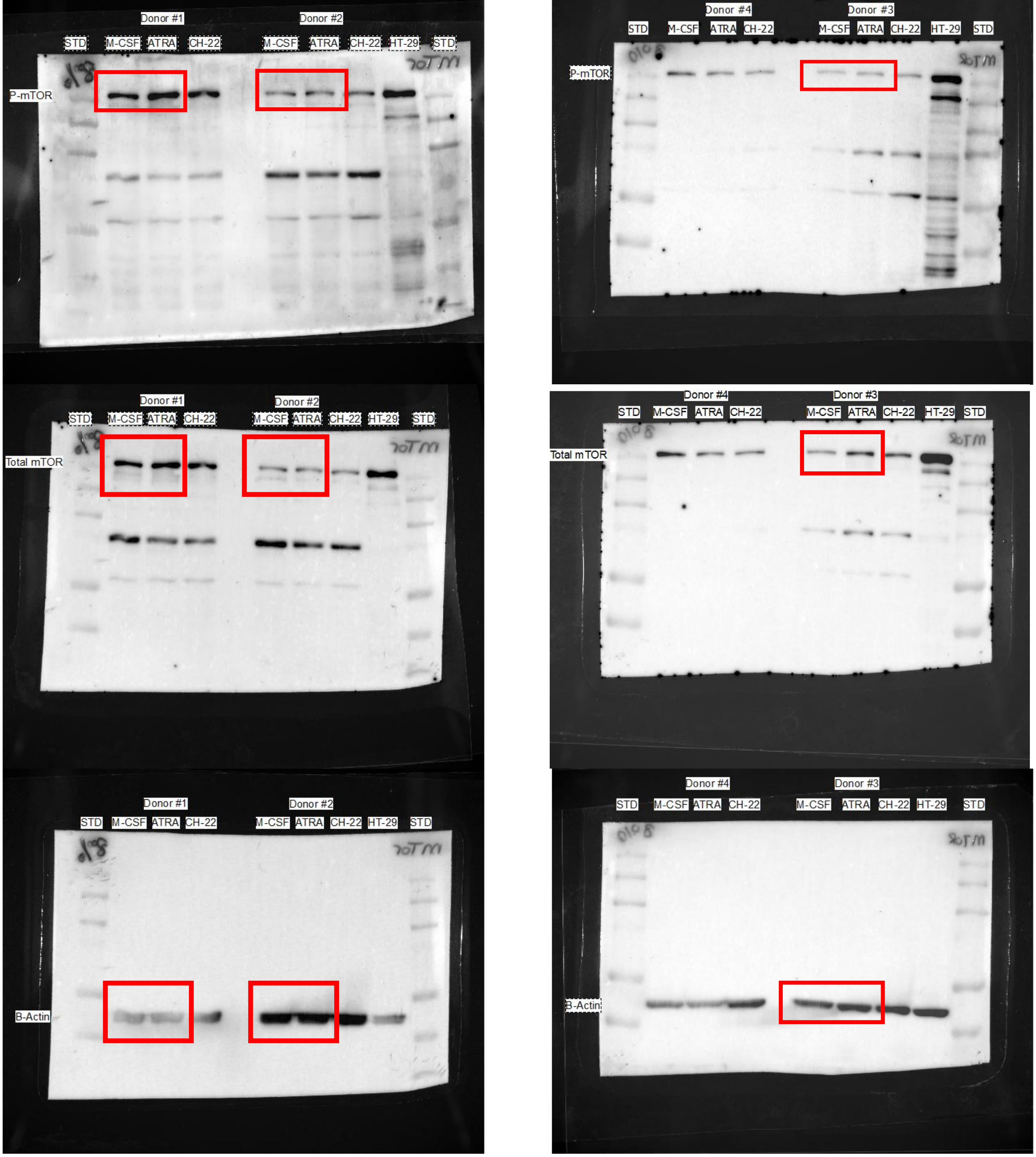


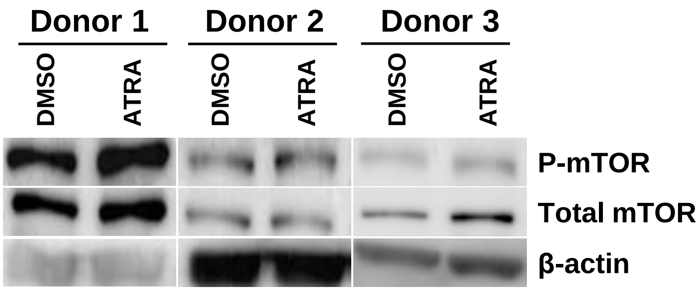


| | Donor 1 DMSO | Donor 1 ATRA | Donor 2 DMSO | Donor 2 ATRA | Donor 3 DMSO | Donor 3 ATRA |
|---|---|---|---|---|---|---|
| **Ratio P/β** | 0.409 | 0.916 | 0.115 | 0.197 | 0,256 | 0.371 |
| **Ratio T/β** | 0.955 | 1.730 | 0.238 | 0.308 | 0.430 | 0.765 |

Dias *et al.*, Supplemental Figure 6A

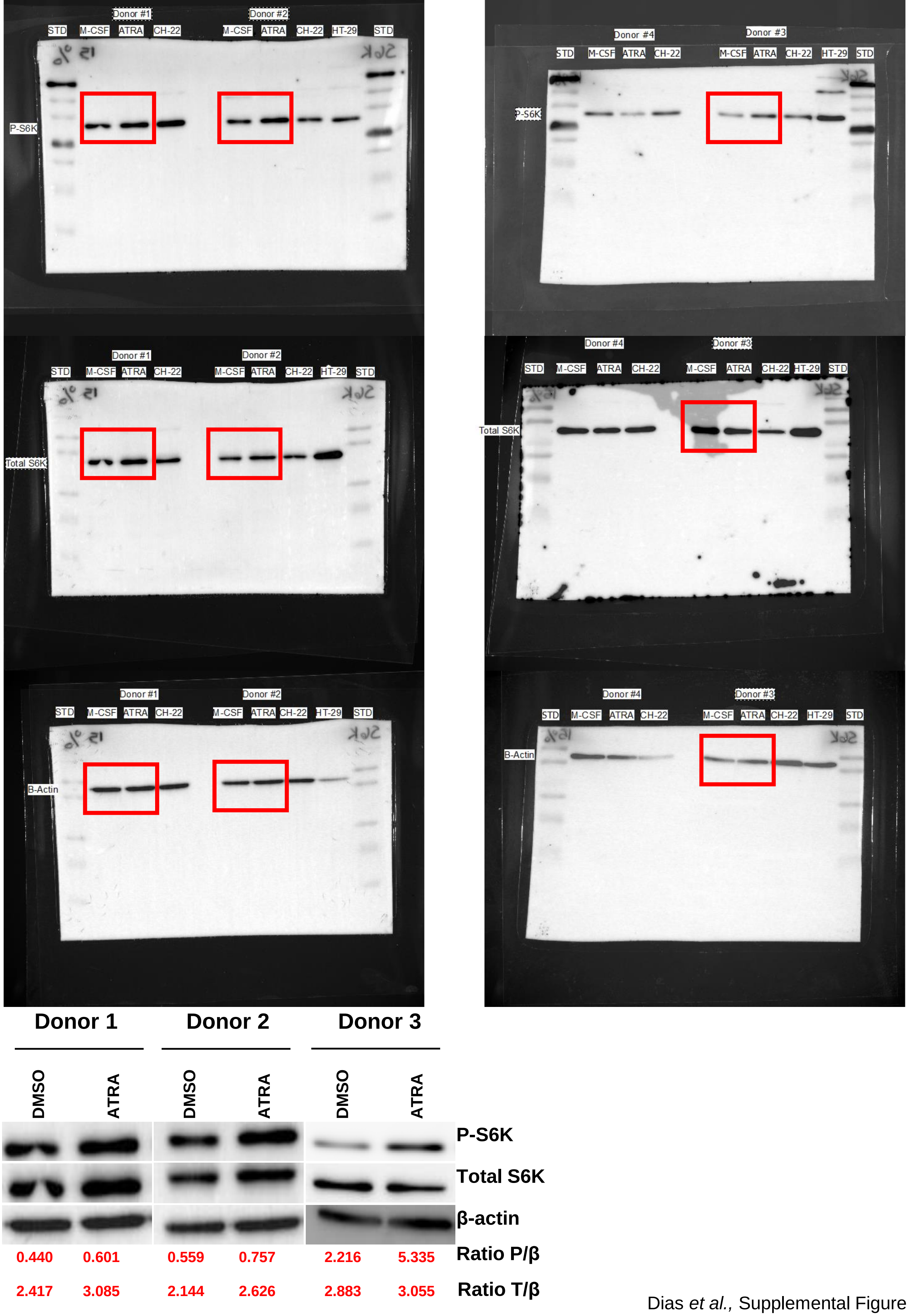


Dias *et al.*, Supplemental Figure 6C

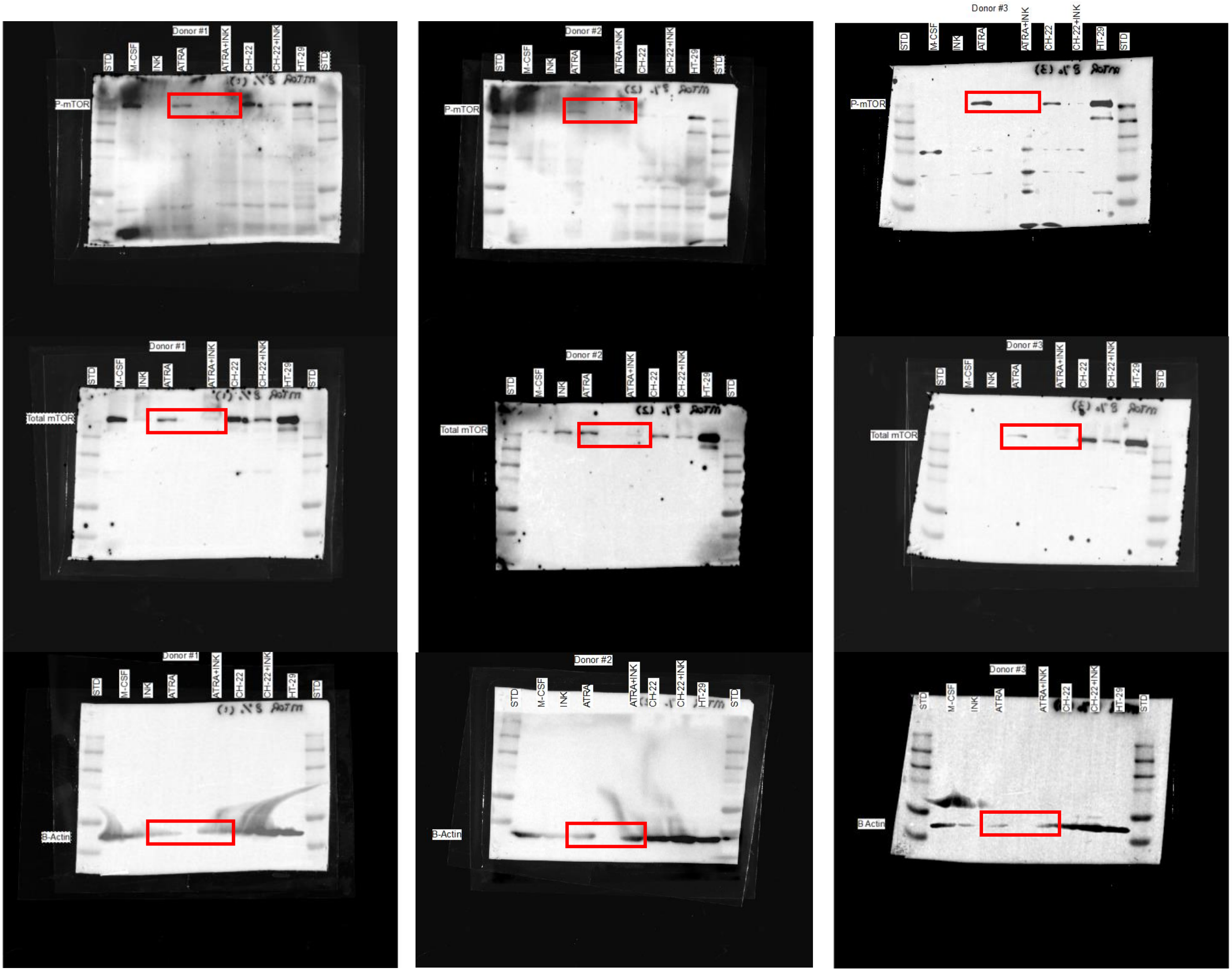


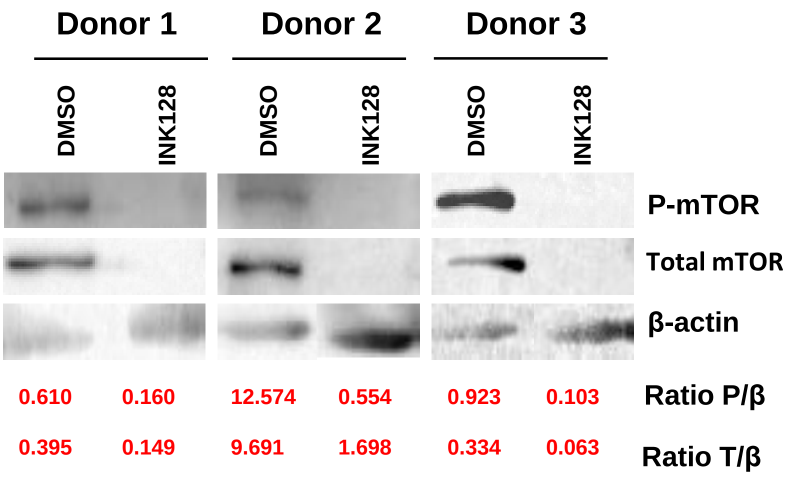


Dias *et al.*, Supplemental Figure 7A

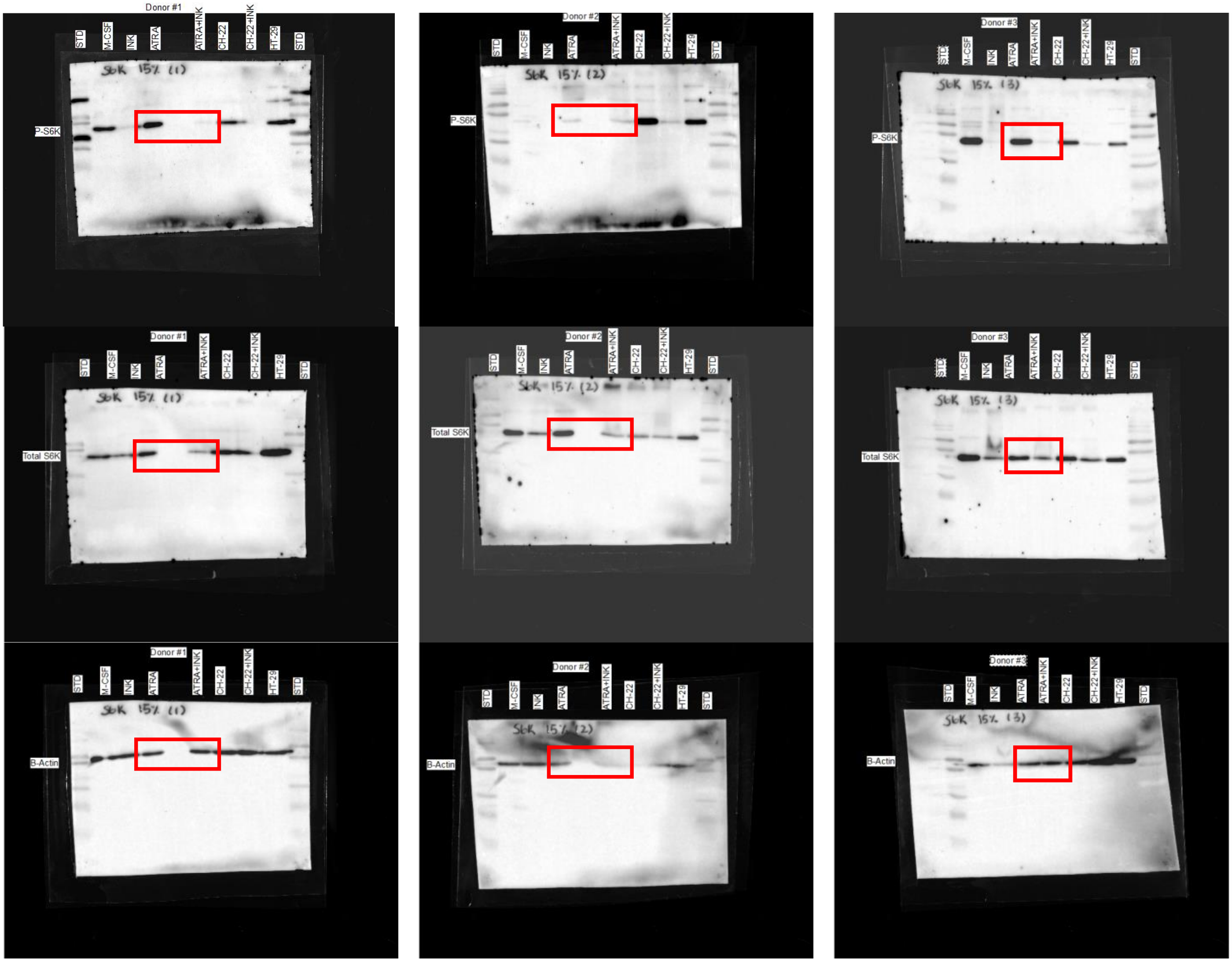


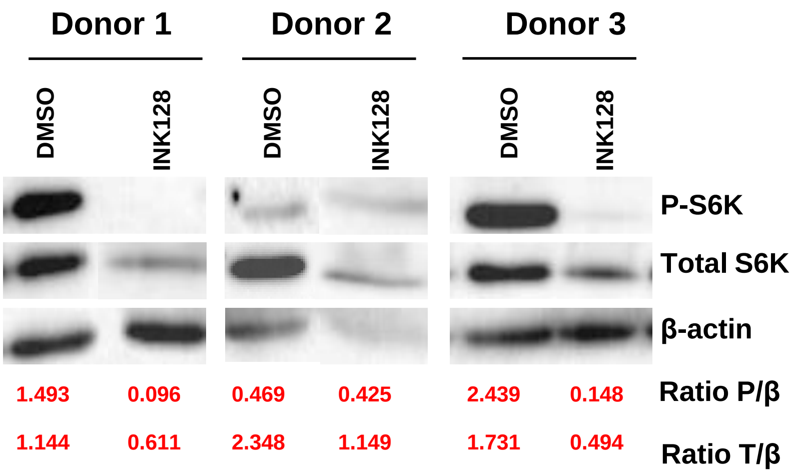


Dias *et al.*, Supplemental Figure 7C

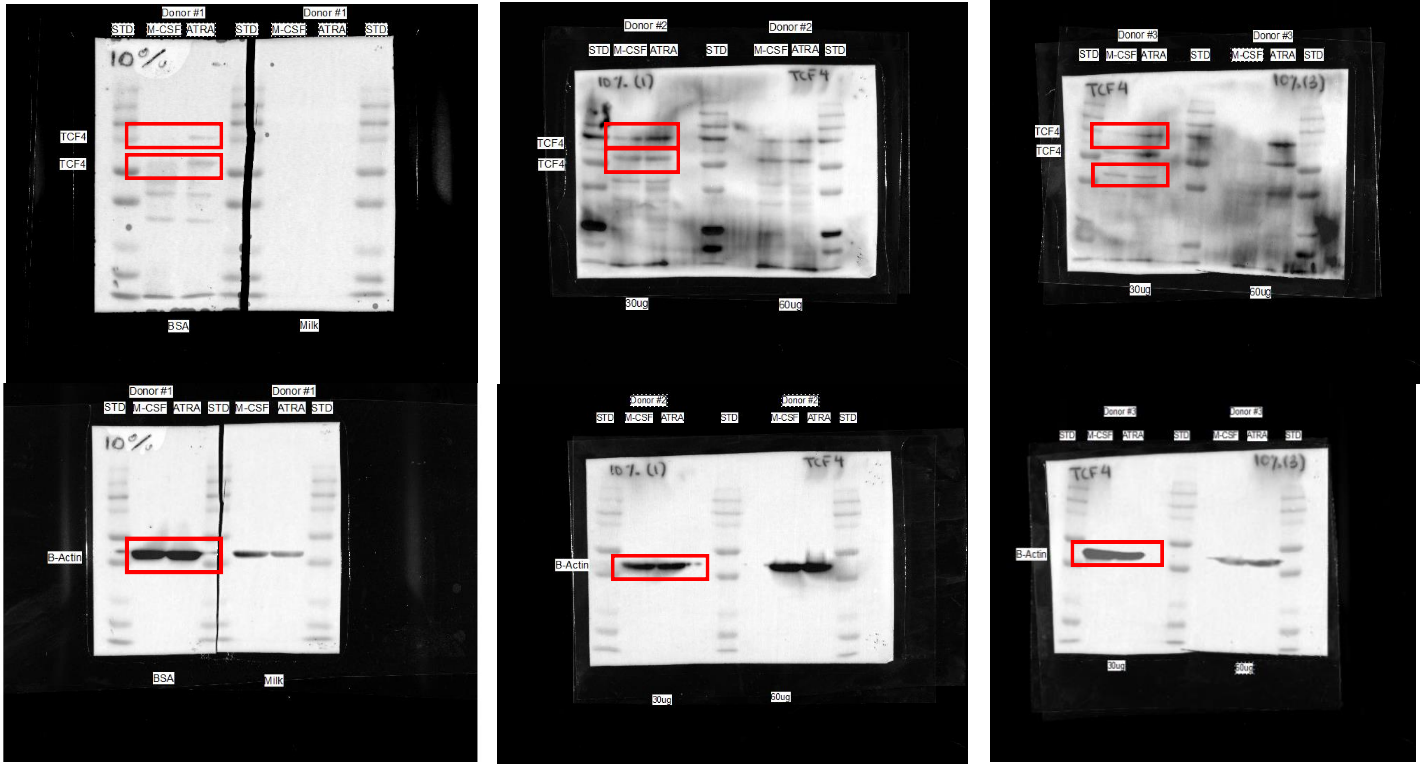
Donor #1
STD M-CSF ATRA STD M-CSF ATRA STD
TCF4
TCF4
BSA
Milk
Donor #2
STD M-CSF ATRA STD M-CSF ATRA STD
30ug
60ug
Donor #3
B-Actin

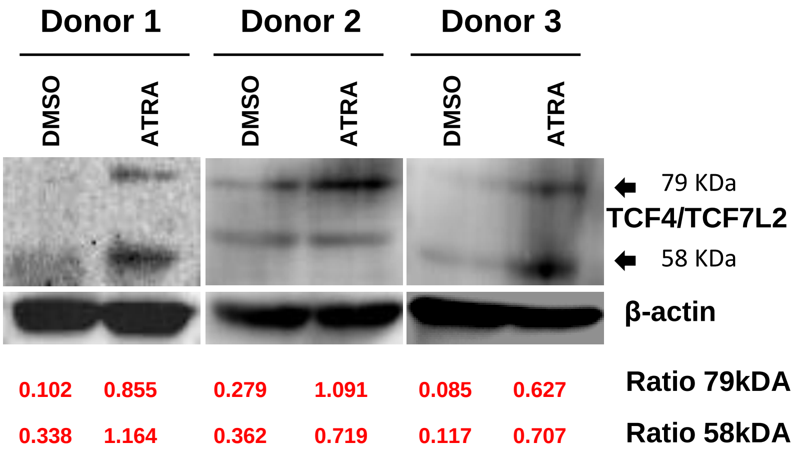
Donor 1
Donor 2
Donor 3
DMSO
ATRA
DMSO
ATRA
DMSO
ATRA
79 KDa
TCF4/TCF7L2
58 KDa
β-actin
0.102 0.855 0.279 1.091 0.085 0.627
Ratio 79kDA
0.338 1.164 0.362 0.719 0.117 0.707
Ratio 58kDA
